# Single-cell RNA-sequencing data analysis reveals a highly correlated triphasic transcriptional response to SARS-CoV-2 infection

**DOI:** 10.1101/2022.06.03.494642

**Authors:** Pablo A. Gutiérrez, Santiago F. Elena

## Abstract

Single-cells RNA sequencing (scRNA-seq) is currently one of the most powerful techniques available to study the transcriptional response of cells to external perturbations. However, the use of conventional bulked RNA-seq analysis methods can miss important patterns underlying in the scRNA-seq data. Here, we present a reanalysis of scRNA-seq data from human bronchial epithelial cells and colon and ileum organoids using pseudo-time profiles based on the degree of virus accumulation which reflect the progress of infection. Our analysis revealed a transcriptional response to infection characterized by three distinct up- and down-regulatory phases, that cannot be detected using classical two-group comparisons. Interrogation of results, focused on genes involved in interferon-response, transcription factors and RNA-binding proteins, suggests a highly correlated transcriptional response for most genes. In addition, correlation network analysis revealed a distinct response of genes involved in translation and mitochondrially-encoded genes. Based on our data, we propose a model where modulation of nucleocytoplasmic traffic by the viral protein nsp1 explains the triphasic transcriptional response to SARS-CoV-2 infection.

## INTRODUCTION

Since the beginning of the SARS-CoV-2 pandemic in 2019, an unprecedented global research effort has taken place to elucidate the detailed molecular biology of this virus and its effects within infected cells (Hu et al., 2020; Barrantes, 2021; Brant et al., 2021). Among the vast amount of information generated, single-cell RNA sequencing (scRNA-seq) datasets are particularly interesting as they can provide a detailed picture of the transcriptional state of individual cells at different stages during the infection cycle (Ciuffi et al., 2016; Cristinelli & Ciuffi, 2018). This is possible because the synthesis of cDNA libraries is performed on individualized cells and each cDNA product is labelled with an identifier that provides unique marker for of each transcribed molecule (Hwang et al., 2018; Ziegenhain et al., 2017; Ding et al., 2020). Therefore, instead of describing the average transcriptional state of thousands to millions of cells, as in bulked RNA-seq, scRNA-seq captures a continuum of transcriptional states that can be used as proxy to reconstruct the chronological response of a cell population to stimuli (Chen et al., 2019; Kharchenko, 2021; Ding et al., 2022).

In virology, scRNA-seq was first used to study the heterogeneity of hepatitis C virus quasiespecies in liver cells (McWilliam & McLauchlan, 2013). Since then, most virology scRNA-seq studies have focused on characterizing the heterogeneity of host cells infected with a wide range of viruses such as human papilloma virus, West Nile virus, yellow fever virus, Zika virus, or human immunodeficiency virus type 1, among others (Cristinelli & Ciuffi, 2018; Russell et al., 2018; Steuerman et al., 2018). In addition, more recent studies have started to address questions related to the transcriptional dynamics of viruses and infected cells upon infection (Ciuffi et al., 2016; Shnayder et al., 2018; Zanini et al., 2018). SARS-CoV-2 is no exception, and since the emergence of the COVID-19 pandemic, dozens of papers have been published addressing different aspects of the molecular biology of this virus (Ziegler et al., 2020; Ramirez et al., 2021; Schuler et al., 2021; Sen et al., 2021). Unfortunately, despite the large volume of available data, most scRNA-seq investigations on SARS-CoV-2 have focused on characterizing cell tropisms or molecular mechanisms at macroscopic timepoints along the infection cycle.

In this study, we present an analysis of the host cell response during the progression of SARS-CoV-2 infection using virus accumulation as a proxy of time. To obtain common features of the SARS-CoV-2 response, we performed a meta-analysis of three publicly available scRNA-seq datasets derived from *in vitro* inoculation studies of human bronchial epithelial cells (Ravindra et al., 2021), and colon and ileum organoids (Triana et al., 2021). Our results revealed that the average transcriptional gene response to SARS-CoV-2 infection is non-linear and cannot be described adequately using conventional bulked RNA-seq analysis methods. Instead of conventional two-group comparisons (*i.e*., infected *vs* noninfected), our analyses of the datasets involved pairwise comparisons of whole transcriptional profiles to average transcriptional responses to within-cell virus accumulation. Our results revealed that about 90% of genes exhibited the same qualitative response to infection suggesting that transcription of most genes is modulated by the same mechanism. Additionally, a strong correlated response was found among mitochondrial encoded genes and genes involved in protein translation. A mechanism explaining the genome-wide transcriptional response to SARS-CoV-2 infection is proposed.

## MATERIALS AND METHODS

### Data

Raw sequencing files were downloaded from the Sequence Read Archive (SRA) at NCBI using the SRA toolkit 2.11.1 (https://www.ncbi.nlm.nih.gov/sra/) and included previously published scRNA-seq data on SARS-CoV-2 infection of human bronchial epithelial cells (hBECs) (Ravindra et al., 2021) and human intestinal epithelial cells (hIECs) (Triana et al., 2021). The hBECs data comprised transcripts from a mock infection, and cells at 1-, 2-, and 3-days post-infection (dpi) with SARS-CoV-2 isolate USA-WA1 /2020 (Ravindra et al., 2021). The hIECs data was produced from colon- and ileum-derived organoids infected with SARS-CoV-2 isolate strain BavPat1 (Triana et al., 2021), and comprised transcripts from a mock infection, and infected cells at 12- and 24-hours post-infection (hpi). In both studies, gel bead-in emulsions (GEMs) were prepared from single-cell suspensions using the 10× Genomics Single Cell 3’ Library Kit NextGem V3.1 (10× Genomics, USA). hBECs libraries were sequenced on a NovaSeq600 system (Ravindran et al., 2021) and hIECs using a HiSeq4000 system (Triana et al., 2021).

### Data processing

Datasets were processed using custom Python scripts. Only sequences with a perfect match to the 10× genomics whitelist (3M-february-2018.txt.gz) and no ambiguous nucleotide calls in the UMI portion were used in the analysis. Selected sequences were mapped to a custom database comprising the complete set of reference human messenger RNAs available at NCBI, mitochondrial-encoded transcripts from the human reference mitochondrial genome (NC_012920), and SARS-CoV-2 genomes corresponding to the reference sequence Wuhan-Hu-1 (NC_045512), and isolates USA-WA1 /2020 (MW811435) and BavPat1 (MZ558051) used for infection in both datasets (Supplementary Data 1). Read mapping was performed with MagicBLAST (Boratyn et al., 2019). Only sequences that mapped unambiguously to a single gene with more than 95% coverage and a single UMI were included in this analysis. In contrast to the original published analyses (Ravindra et al., 2021, Triana et al., 2021), we selected GEMs with respect to the total number of transcripts (UMIs) instead of the total number of transcribed genes. Empty GEMs and multiplets were removed from the datasets using a custom Python script that ordered GEMs with respect to the number of transcripts transformed using a log_10_ scale; then local standard deviation of the ordered data was calculated using window size of five datapoints. Upper and lower thresholds delimiting multiplets and void beads were determined using the 1.5×IQR rule (Table 1, Supplementary Fig. 1). Mapping information from the selected cells was then transformed into a matrix of counts which included genes detected in at least one hundred cells (Supplementary Data 1). Furthermore, in contrast to the original analysis of these datasets, we did not discard cells with a high percentage of transcribed mitochondrial genes as overexpression of mitochondrial genes could be a legitimate response to SARS-CoV-2 infection (Ilicic et al., 2016; Miller et al., 2021).

### Estimation of gene frequencies

The probability of detecting *n* transcripts from a gene in a cell depends on its relative frequency and the total number of sampled transcripts. Therefore, as cDNA amplification of cellular transcripts represent only 10 to 20% of the total mRNA content (Islam et al., 2014; Rostom et al., 2017; Hwang et al., 2018), gene count data from scRNA-seq contains many missing values, or dropouts, that result from Poisson sampling of low abundance transcripts in cells with few sequenced transcripts (Svensson et al., 2017; Chen et al., 2019). The probability of observing *n_gi_* read counts for a given gene *g* ∈ {1, 2, …, *G*} across cells *i* ∈ {1, 2, …, *I*} can be modelled using a Poisson distribution were the likelihood of observing *n_gi_* counts given a particular transcript abundance *θ _gi_* is given by:

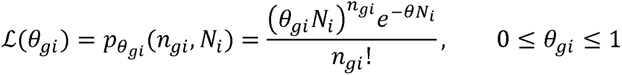

with 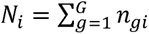.

In this work gene abundances were calculated as the weighted average of gene frequencies (*θ*) with respect to the normalized likelihood function 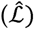 and multiplied by a scaling factor of 10^4^ (transcripts per ten thousand or *TPT*):

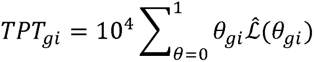

with 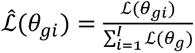.

An uncorrected normalized matrix was also calculated. Corrected and uncorrected frequency matrices are available in Supplementary Data 2.

### Volcano plot analysis

Basal levels of expression for each gene were determined using the average expected *TPT* values from mock cells with at least one gene count per gene of interest. Infected cells, on the other hand, were binned with respect to viral loads using a log_2_(*TPT*_SARS-CoV-2_ + 1) scale. Only cells with at least one count for SARS-CoV-2 and the target gene were used. *P*-values were calculated using a two-tailed Mann-Whitney *U* test and corrected using the Benjamini-Hochberg method (Benjamini & Hochberg, 1995) using a significance threshold of 0.01. Differential expression analysis results are provided in Supplementary Data 3.

### *Z*-score profiles and analysis

Normalized transcriptional responses were calculated using a z-score of the transcription levels observed at each log_2_(*TPT*_SARS-CoV-2_ + 1) corrected by the levels and standard deviation observed in the uninfected cells. For each cell type, a mean response vector was calculated by averaging the normalized response of all genes. Pairwise comparisons between individual transcriptional profiles and the mean response vector were performed using a root-mean-square deviation (*RMSD*) metric and the magnitude of the initial response (Δ_0_) measured as the average z-score of the first five datapoints. A transcriptional response was considered an outlier when either the *RMSD* or Δ_0_ was outside the intervals defined by the 1.5×IQR rule.

### Gene Ontology (GO) analysis

Analysis of biological functions was performed using a custom database of high-quality GO terms of human proteins downloaded from the Uniprot databaset (UniProt Consortium, 2015; Supplementary Data 3). Quantification of the most abundant GO terms was performed using custom scripts that mapped selected genes to a dictionary of GO annotations. GO enrichment analysis was performed by comparison of gene subsets to the list of mapped genes as background. *P*-values were calculated using the Fisher’s exact test and corrected Benjamini-Hochberg method (with threshold significance of 0.01).

### Construction of gene regulatory networks

Undirected correlation networks were built using the normalized transcription profiles. A link between two genes was stablished when the two sample Kolmogorov-Smirnov test indicated that both *z*-score distributions followed the same distribution (*P* ≤ 0.01) and the corrected Pearson correlation coefficient between expression profiles was *r* ≥ 0.9. In both instances, *P*-values were corrected using the Benjamini-Hochberg method (Benjamini & Hochberg, 1995). Only those genes which had at least 50 measurements in common were included in the analysis (Supplementary Data 5). Network properties were characterized with the Python networkx package (Aric et al., 2008) and visualized with Gephi (Bastian et al., 2009).

### Data and Code Availability

All data generated or analyzed during this study are included in this article and its supplementary information files. Tables and processed datasets are provided as <u>supplementary data</u>. Codes that support the findings of this research are available at GitHub (https://github.com/paguties/SARS-CoV-2_scRNAseq.git).

## RESULTS

### The global transcriptional response to SARS-CoV-2 exhibits a wave-like pattern

First, we tried to understand the global transcriptional response of infected cells by plotting the distribution of log_2_ fold-changes (log_2_*FC*) of transcripts as a function of the log_2_ of SARS-CoV-2 levels (Fig. 1a, Supplementary Data 1). This analysis indicated that the average transcriptional response displays an oscillating behavior that can be divided qualitative into three phases. An early phase that occurs at low viral RNA levels (≲ 1 *TPT*) and is characterized by a global shutdown of transcription were 99.5% of genes in hBECs, 98.7% in colon and 98.0% in ileum are downregulated. An intermediate phase takes place at intermediate viral loads and is characterized by an increase in expression with maximum fold changes of between 1.4 and 4.1 in ileum. Finally, the late phase was observed in cells where SARS-CoV-2 transcripts dominate the transcriptome and cellular transcripts are, again, significantly below the levels of uninfected cells.

**Figure 1.**
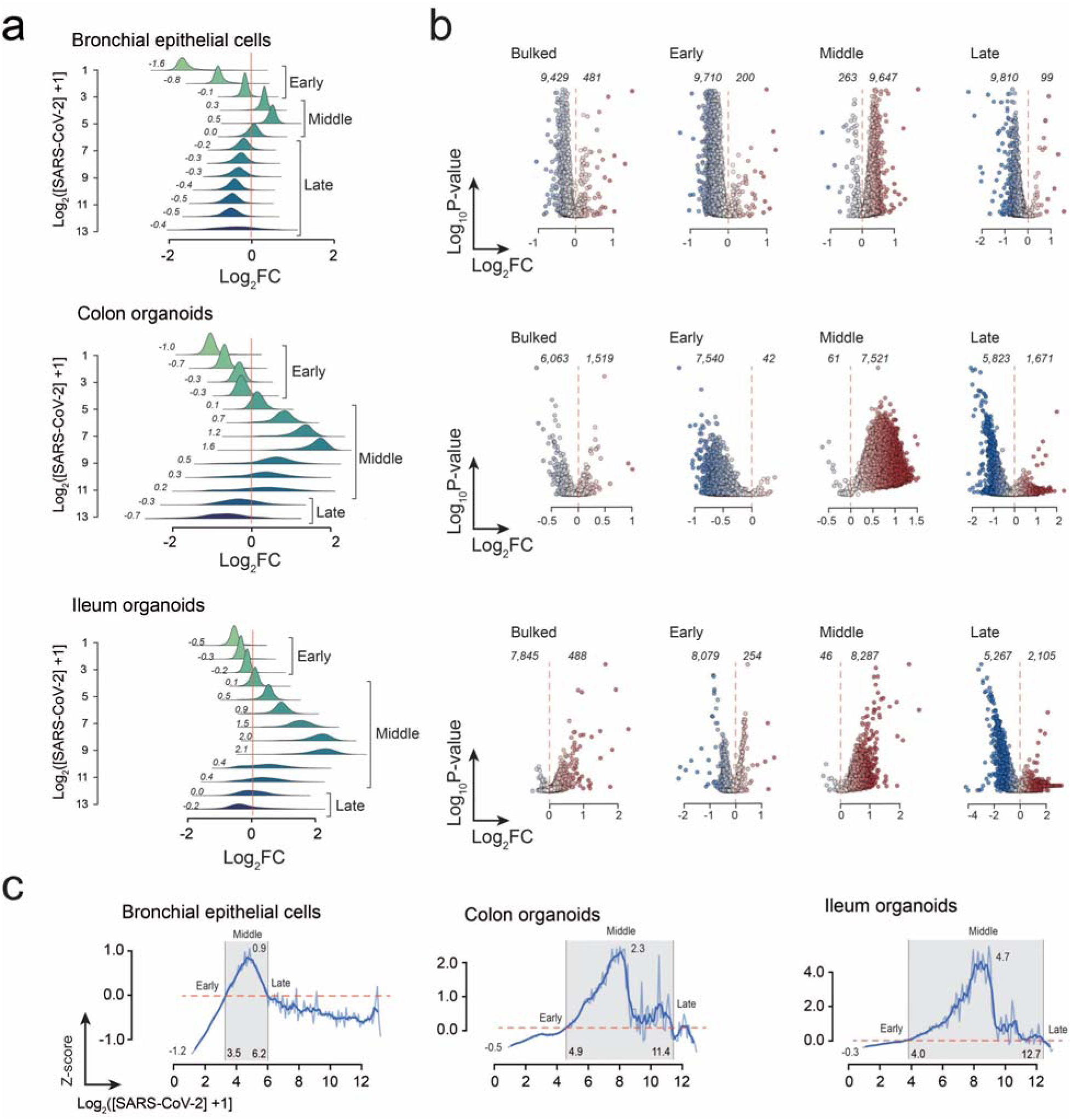
Global transcriptional response to SARS-CoV-2 in human bronchial epithelial cells, colon- and ileum-organoids. **a** Distribution of log_2_*FC* values for all genes plotted at log_2_[SARS-CoV-2 +1] intervals. Mean log_2_*FC* values are shown to the left to each histogram. **b** Volcano plot analysis of infected cells with respect to non-infected cells using bulked cells, and subgroups comprising cells at early, middle, and intermediate phases of infection. For clarity, corrected *P*-values were removed from the ordinate axis. **c** The average *z*-score plot confirms the global oscillatory transcriptional profile of genes at increasing viral loads.

As the oscillatory transcriptional response to SARS-CoV-2 might bias conventional bulked differential gene expression (DEG) analyses, we tested the effect of selecting cells from different phases as contrasting groups. In Fig. 1b, we show the volcano plots that result from comparison with all the infected cells, and with three subsets comprising early, intermediate, and late phase cells (Fig. 1b). These results unveil that the classification of genes as down- or up-regulated is highly dependent on how the contrasting cell populations are selected. For example, in hBECs, the bulked DEGs analysis revealed 9,429 down- and 481 up-regulated genes; and similar results obtained using cells from the early phase (9,710 up- *vs* 200 down-regulated genes). However, this trend is reversed when cells from the intermediate phase are used: 9,647 up- *vs* 263 down- regulated genes are observed. Finally, as expected from the distribution of log_2_*FC* values, DEG trends are reversed again when cells from the late phase are compared (99 up- *vs* 8,810 down- regulated genes). Similar results were obtained in the analysis of infected colon and ileum cells (Fig. 1b, Supplementary Data 3).

To better compare the global transcriptional response at different stages of the viral infection cycle, we plotted the average scaled gene expression values at different viral loads (Fig. 1c). To do this, expression values at each log_2_ interval were transformed into *z* scores with respect to the mean and standard deviations of uninfected cells. Under this transformation, up- and down- regulated regimes correspond to positive and negative values, respectively. Data ordered in this way can be interpreted as a pseudotime, which, here represents doublings in viral concentration (Griffiths et al., 2018; Chen et al., 2019, Ding et al., 2021). In hBECs, *z*-scores fluctuated between −1.2 and 0.9, and transitions between phases occurred at pseudotimes of 3.5 (∼10.3 *TPT*) and 6.2 (∼72.5 *TPT*) (Fig. 1c). In colon- and ileum-organoids down-regulation during the early phase was less pronounced but exhibited a stronger up-regulatory response in the intermediate phase. In these cases, *z*-scores fluctuated between −0.5 and 2.3 in colon, and −0.3 and 4.7 in ileum. Transition points occurred at pseudo times of 4.9 (∼28.9 *TPT*) and 11.4 (∼2,701.3 *TPT*) in colon, and 4.0 (∼15 *TPT*) and 12.7 (∼6,653 *TPT*) in ileum (Fig. 1c).

### Classification of gene responses

Next, we proceeded to identify genes with abnormal transcriptional profiles using two metrics: the root-mean-square deviation (*RMSD*) to the average response vector and the magnitude of the initial response (Δ_0_) (Fig. 2a). The latter was used to give a stronger weight to data points at earlier infection times and to select for genes where the initial transcriptional response was weaker or stronger. A transcriptional profile was considered an outlier when either of these parameters was outside the intervals defined by the 1.5×IQR rule. Our results indicate that most cellular transcripts have a response pattern qualitatively similar to the mean response vector, with only 7.2% to 10.1% of genes exhibiting atypical transcriptional profiles (Fig. 2b, Supplementary Data 3); 183 outliers were shared between the three cell types (Fig. 2c). A gene ontology (GO) analysis of all the outliers from the three cell types (Fig. 2d) revealed that this subset was enriched in genes involved in immune response (*e.g*., GO:0030593, GO:0031640 and GO:0061844), translation (*e.g*., GO:0042274, GO:0000028, GO:000641 and GO:0002181), respiration (*e.g*., GO:0006120 and GO:0042776), and control of cellular proliferation and death (*e.g*., GO:0001895). The subset was also significantly depleted in genes involved in intracellular protein transport (*e.g*., GO:0006886), protein phosphorylation (*e.g*., GO:0006468) and regulation of transcription (*e.g*., GO:0006355 and GO:0006357).

**Figure 2.**
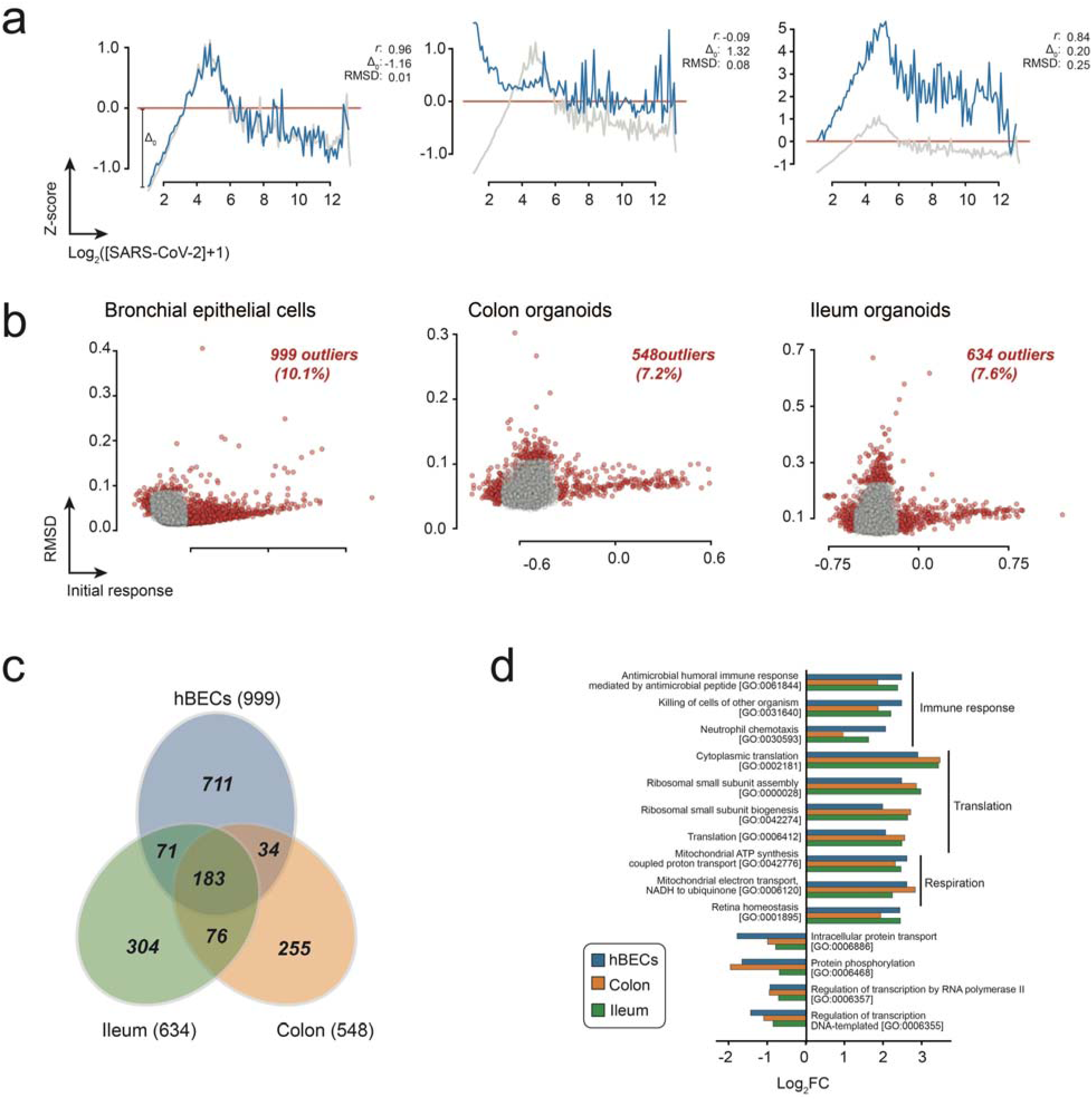
Classification of genes based on their similarity to the mean response vector. **a** Examples of selected individual transcriptional profiles. Each individual transcriptional profile (blue) was compared to the mean response vector (gray) using four parameters: the Pearson correlation (*r)*, the root-mean-square-deviation (*RMSD*) and the magnitude of the initial response (Δ_0_). **b** Classification of genes as normal responding (gray) and outliers (red) using *RMSD* and Δ_0_. **c** Venn diagrams illustrating the distribution of outlier among the three datasets used in the analysis. **d** GO-enrichment analysis of outliers from the three datasets.

### Transcriptional response of IFN-stimulated genes

At early infection times SARS-CoV-2 downregulates the expression of IFN-stimulated genes (ISGs) involved in antiviral defense (Schoggins et al., 2011; Thoms et al., 2020; Finkel et al., 2021). We then decided to investigate if the IFN response follows an abnormal transcription pattern upon SARS-CoV-2 infection. Firstly, we analyzed the transcriptional profiles of genes encoding for MDA-5 and RIG-I which are the main intracellular sensors that recognize double-stranded RNAs and trigger the innate immune response upon infection (Rebendenne et al., 2021; Sampaio et al., 2021; Thorne et al., 2021). Analysis of the transcriptional profiles of MDA-5 and RIG-I, encoded by genes *IFIH1* and *DDX58*, respectively, revealed that both have a transcriptional response like the average response vector (Fig. 3a upper panels, Supplementary Data 3). This result suggests that the transcriptional shutdown of the IFN response induced by MDA-5 and RIG-I is probably not specific, and results from the global transcriptional shutdown that affects most genes in the infected cell. A GO enrichment analysis of terms associated with the MDA-5 and RIG-I signaling pathways revealed a progressive increase in the proportion of genes involved in these pathways that reaches a plateau during the intermediate infection phase (Fig. 3a, lower panels), which agrees with experimental evidence suggesting a delayed response of the IFN response in infected cells (Banerjee et al., 2020; Xia et al., 2020; Kumar et al., 2021; Thorne et al., 2021).

**Figure 3.**
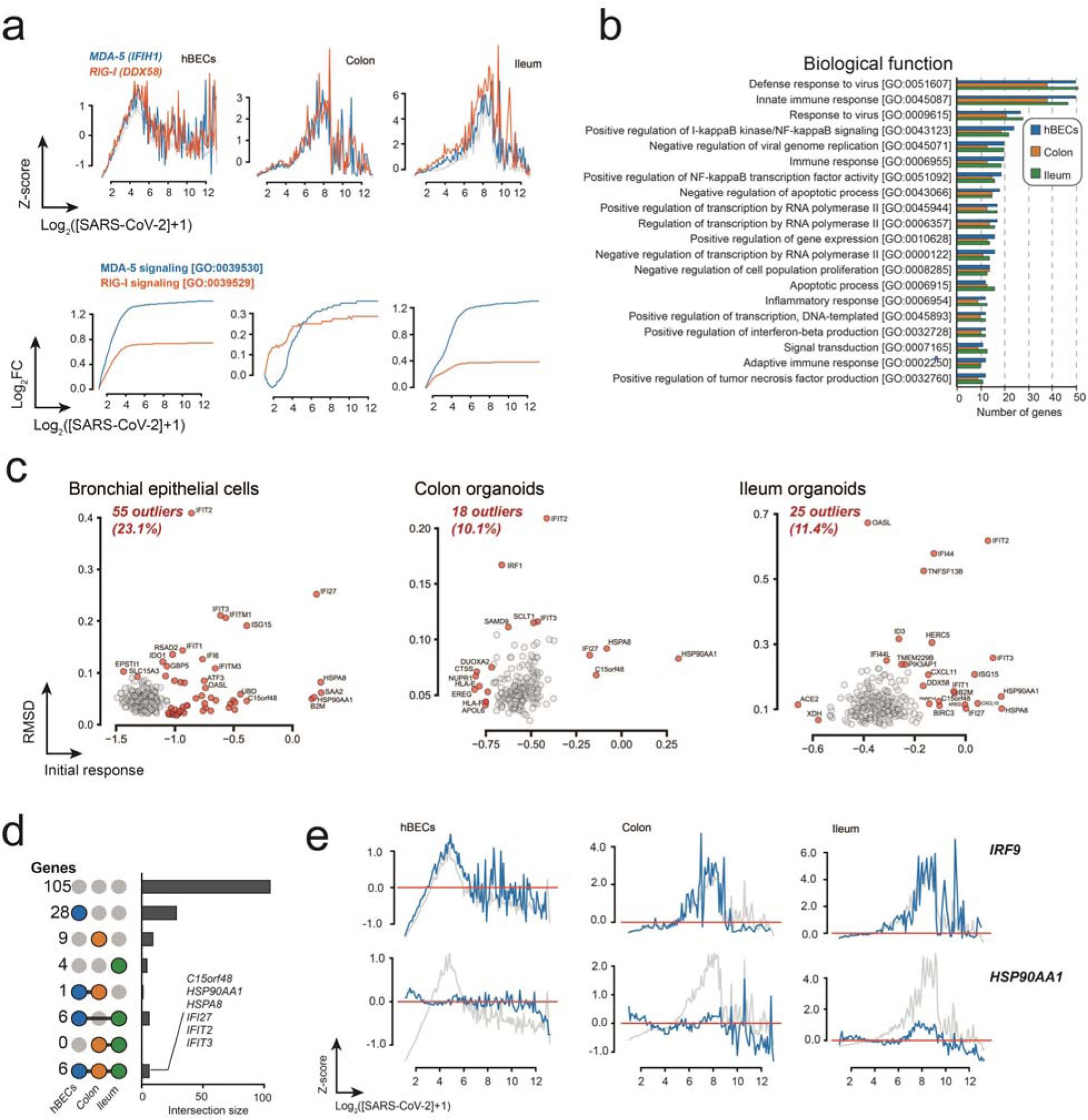
Global transcriptional response of ISGs in human bronchial epithelial cells, colon- and ileum-organoids. **a** The transcriptional response of the cytoplasmic viral receptors MDA-4 and RIG-I correlated with the mean transcriptional response. Enrichment of GO term associated with both signaling pathways are shown below. **b** GO annotation of the 20 most common terms in the identified ISGs. **c** Distribution of *RMSD* and Δ_0_ profiles of outlier ISGs. **d** Upset plots showing the total the classification of genes between common ISGs. Grays correspond to genes with average transcriptional response, black represents genes with abnormal transcriptional responses. **e** Representative examples of an ISG with average (*IRF9*) and abnormal (*HSP90AA1*) transcriptional responses.

We then proceeded to analyze the response of individual ISGs in the three datasets. To do this, we selected genes using a recently curated and validated dataset of ISGs (Martin-Sancho et al., 2021). A total of 238, 179, and 219 ISGs were identified in hBECs, colon and ileum, respectively; with 159 of these genes in common among the three datasets (Fig. 3a, Supplementary Data 3). As expected, a GO analysis revealed that most of the identified ISGs are involved in the defense response to viruses (Fig. 3b). Like the MDA-5 and RIG-I receptors, 77% - 90% of ISGs exhibited an average transcriptional response to SARS-CoV-2 indicating that the mRNA levels of these genes are negatively regulated at the onset of infection and their response is probably part of the global transcriptional shutdown (Fig. 3). An analysis of abnormal transcriptional responses identified 55 genes from hBECs, 18 from colon and 25 from ileum; six of them common to the three datasets: heat shock protein HSP90α (*HSP90AA1*), interferon-induced protein with tetratricopeptide repeats 2 (*IFIT2*), interferon-induced protein with tetratricopeptide repeats 3 (*IFIT3*), heat shock cognate 71 kDa protein (*HSPA8*), normal mucosa of esophagus-specific gene 1 protein (*C15orf48*), and interferon alpha-inducible protein 27 (*IFI27*) (Fig. 3c). HSP90AA1 and HSPA8 are chaperone proteins which are typically required for viral replication (Geller et al., 2012). In addition, HSPA8 has been shown to be an attachment factor for the avian infectious bronchitis coronavirus (Zhu et al., 2020). *C15orf48* expression, on the other hand, has been shown to be involved in the mitochondrial stress response (MISTR) and is expressed by pathogenic macrophages in severe COVID-19 (Liao et al., 2020; Clayton et al., 2021). Interestingly, IFI27 and IFIT3 are also involved with the mitochondrial processes; the former locates in the nuclear inner membrane and is indispensable for mitochondrial function (Jin et al., 2018) and the latter is mediator of the mitochondrial antiviral signaling (MAVS) complex triggered by the MDA-5 and RIG-I signaling pathways (Liu et al., 2011).

### Transcriptional response of transcription factors

To further understand the effect of SARS-CoV-2 on transcription factors (TF), we first analyzed the enrichment of genes with GO terms associated with positive and negative regulation of DNA-templated transcription (Fig. 4a). Qualitatively, our analysis suggests that transcription factors are enriched at higher viral loads peaking at about 2 and 4 viral doublings (Fig. 4a). We then investigated the effect of SARS-CoV-2 on individual transcription factors from a curated collection of known and likely human transcription factors (1,639 proteins) published by Lambert et al. (2018) (available at http://humantfs.ccbr.utoronto.ca/). Our analysis identified a total of 785, 448 and 531 TFs in hBEC, colon and ileum, respectively, with 409 TFs in common to the three datasets (Supplementary Data 3). Again, most transcriptions factors behaved like the mean response vector, and only 36 TFs in hBECs, 16 in colon and 22 in ileum were classified as outliers (Fig. 4b, Supplementary Data 3). Three TFs were found to be differentially regulated in the three datasets: the Y-box binding protein 1 (*YBX1*), and the proto-oncogene TFs *JUN* and *JUND* (Fig. 4c, Supplementary Data 3). *YBX1* encodes a highly conserved cold shock domain protein with DNA and RNA binding properties implicated in the regulation of transcription and translation, pre-mRNA splicing, DNA reparation, mRNA packaging, cell proliferation, stress response, and apoptosis. It has been shown to be a component of messenger ribonucleoprotein (mRNP) complexes with a role in microRNA processing (Lyabin et al., 2014; Nagasu et al., 2018). The product of gene *JUN* is a component of the AP1 TF complex involved in oncogenic transformation (Meng et al., 2011) and is targeted to the nucleolus where it seems to modulate nucleolar architecture and ribosomal RNA accumulation (Miyake & McDermott, 2022). The expression profile of *JUN* stands out as its transcriptional profile suggests transcriptional activation at later infection times and its similar response in the three datasets suggest that this gene might be a marker that induces apoptosis in infected cells (Fig. 4d). *JUND* encodes a JUND proto-oncogene that is a functional component of the AP1 TF complex and has been proposed to protect cells from p53-dependent senescence and apoptosis (Ruiz et al., 2021). Finally, together, these results suggest that transcription of most TFs is downregulated non-specifically and the only common differential responding TFs seem to be involved in processes related to programmed cell death.

**Figure 4.**
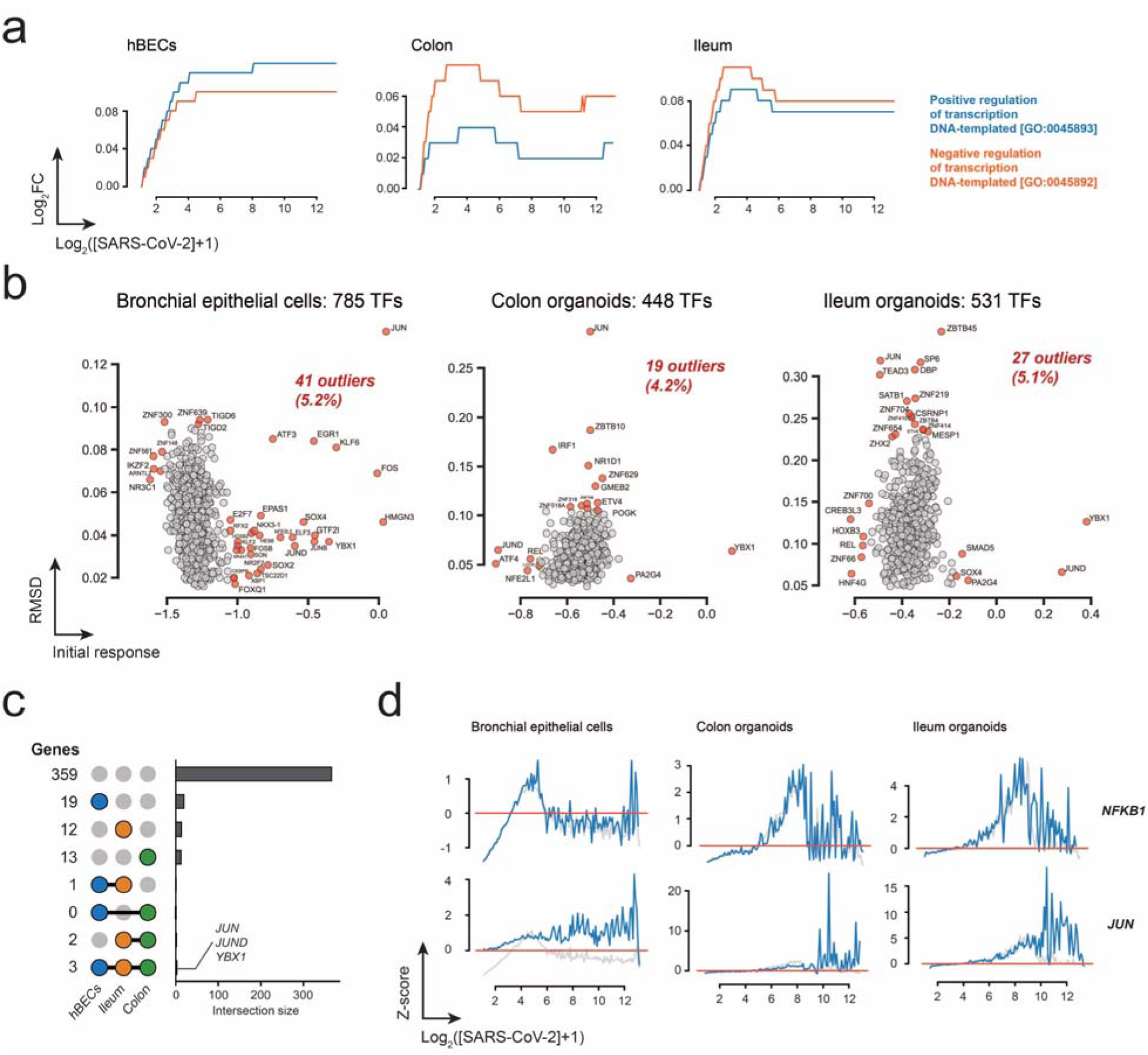
Global transcriptional response transcription factors. **a** Enrichment of GO terms associated with positive and negative regulation of DNA-templated transcription at increasing viral accumulations. **b** Distribution of transcriptional responses of identified transcription factors from hBECs, colon and ileum datasets. Outliers are colored red. c Upset plots showing the total the classification of TFs shared between the three datasets. Grays correspond to genes with average transcriptional response, black represents genes with abnormal transcriptional responses. **d** Representative examples of TFs with average (*NFKB1*) and abnormal (*JUN*) transcriptional responses.

### RNA binding proteins

As levels of mRNA might be modulated by their interaction with RNA binding proteins (RBPs), we explored the transcriptional profiles of genes from a dataset of canonical RBPs from the eukaryotic RBP database, EuRBPDB (Liao et al., 2019). More than a thousand RBPs were identified in each dataset (Fig. 5a, Supplementary Data 3), of which approximately 10% exhibited abnormal transcriptional profiles (Fig. 5b). Interestingly, 83 abnormal responding genes were shared between the three datasets (Fig. 5c) which contrast with the analysis of ISGs and TFs where only a handful of abnormal responding genes shared among datasets. Analysis of genes with an average transcriptional response, revealed that the top GO terms involved functions related to mRNA splicing and mRNA processing (Fig. 5d, upper panel) in agreement with the known disruption of splicing induced by SARS-CoV-2 (Banerjee et al., 2020). In contrast, abnormal responding genes were enriched in terms involved in protein translation (Fig. 5d, lower panel), suggesting a differential response between the RNA processing and protein translation machinery. A detail analysis of the outliers revealed that the common response involved 41 large ribosomal subunit proteins, 30 small ribosomal subunit proteins, four elongation factors (*EEF1A1*, *EEF1B2*, *EEF1G*, and *EEF2*), the FAU ubiquitin-like and ribosomal protein S30 fusion (*FAU*), the histidine triad nucleotide binding protein 1 (*HINT1*), the heterogeneous nuclear ribonucleoprotein A1 (*HNRNPA1*) involved in pre-mRNA processing in the nucleus, nucleolin (*NCL*) involved in ribosome processing, the ras- related nuclear protein (*RAN*) essential for the translocation of RNA and proteins through the nuclear pore complex, the signal recognition particle 14 (*SRP14*) involved in protein targeting to ER, and the ubiquitin A-52 residue ribosomal protein fusion product 1 (*UBA52*) involved in protein degradation by the 26S proteosome, and *YBX1* described above. A STRING analysis of these genes revealed a highly connected physical network that suggest a different regulatory mechanism of genes involved in translation and some of their interacting partners (Fig. 5e).

**Figure 5.**
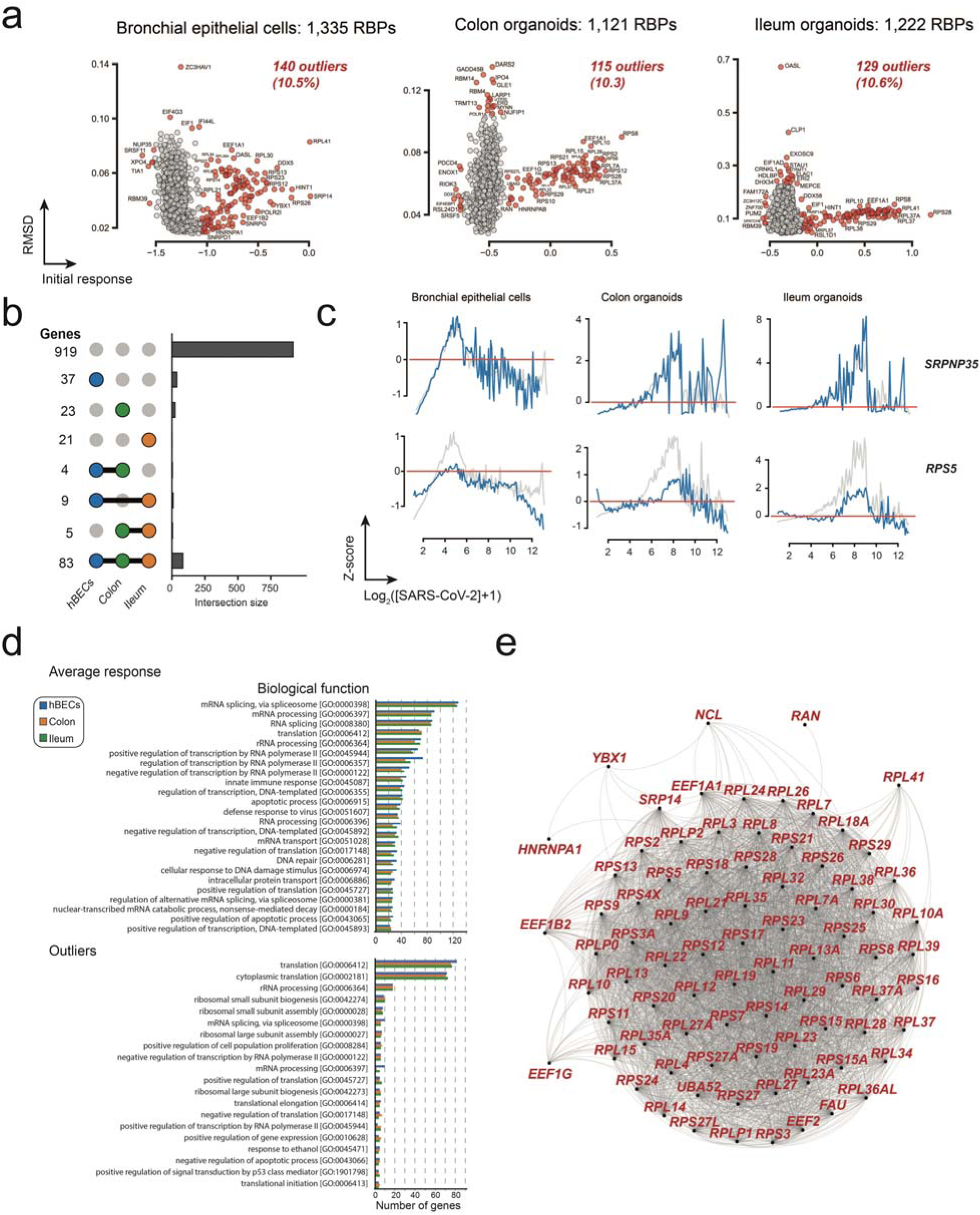
Transcriptional response of RNA binding proteins. **a** Distribution of transcriptional responses of RBPs identified in hBECs, colon, and ileum datasets. Outliers are colored in red. **b** Upset plots showing the total the classification of RBPs shared between the three datasets. Grays correspond to genes with average transcriptional response, black represents genes with abnormal transcriptional responses. **c** Representative examples of RBPs with average (*SRPNP35*) and abnormal (*RPS5*) transcriptional responses. **d** GO annotation of the 25 most common terms of RBPs with normal and abnormal transcriptional responses. **e** Physical network from the STRING database of transcriptional outliers.

### Gene regulatory networks

As gene expression is often modular and usually involves the coordinated expression of functionally related genes (Chen et al., 2019), we investigated the correlation of transcriptional responses between pairs of genes in the three datasets. First, we constructed individual correlation network using the normalized transcriptional profiles (Supplementary Data 2 and 4). Second, we used the individual networks to construct a general response network comprising all the edges shared between the three individual networks (Fig. 6a). The resulting network comprised 25 connected components, the largest of which consisted of 348 genes that exhibited transcriptional profiles similar to the mean response vector (Fig. 6b). The second largest connected component comprised 54 genes, and included mostly ribosomal protein genes, and proteins involved in translation. In this case, all genes within the network exhibited a qualitative behavior that was characterized by weaker downregulation and upregulation of transcription during the early and middle phases of infection as compared to the mean response vector (Fig. 6b). Interestingly, the third largest connected component was comprised exclusively of mitochondrially encoded genes, suggesting that response of mitochondrial genes upon SARS-CoV-2 infection follows a different mechanism from the global response and gene involved in translation (Fig. 6b). The remaining connected components comprised two to three genes and some of them exhibited transcriptional responses similar to those observed in the largest components (Supplementary Data 4); they were no included as part of the largest subnetworks as the some of the edges failed to meet the filtering criteria described. Previous studies have suggested a role of the 5’ UTR of mRNAs in the ability to escape the global suppression of translation induced by SARS-CoV-2 and related coronaviruses (Huang et al., 2011; Nakagawa et al., 2018; Rao et al., 2021). To investigate the presence potential sequence motifs in transcripts from the largest connected components, we generated a sequence logo using the first 20 nucleotides of reference sequences from these clusters. This analysis suggests that transcripts from the first connected component tend to have A, C or G as the first nucleotide with equal probability and higher GC content between positions 7-20 (Fig. 6c). In contrast, sequences from the second connected component contain TOP-motifs in their 5’ UTRs, which is to be expected for transcripts encoding ribosomal proteins and components of the translation apparatus (Meyuhas & Kahan, 2015; Rao et al., 2021). Interestingly, analysis of protein translation in cells expressing SARS-CoV-2 protein Nsp1 revealed that the translation efficiency was slightly higher for genes with lower GC content in their 5’ UTR (Rao et al., 2021).

**Figure 6.**
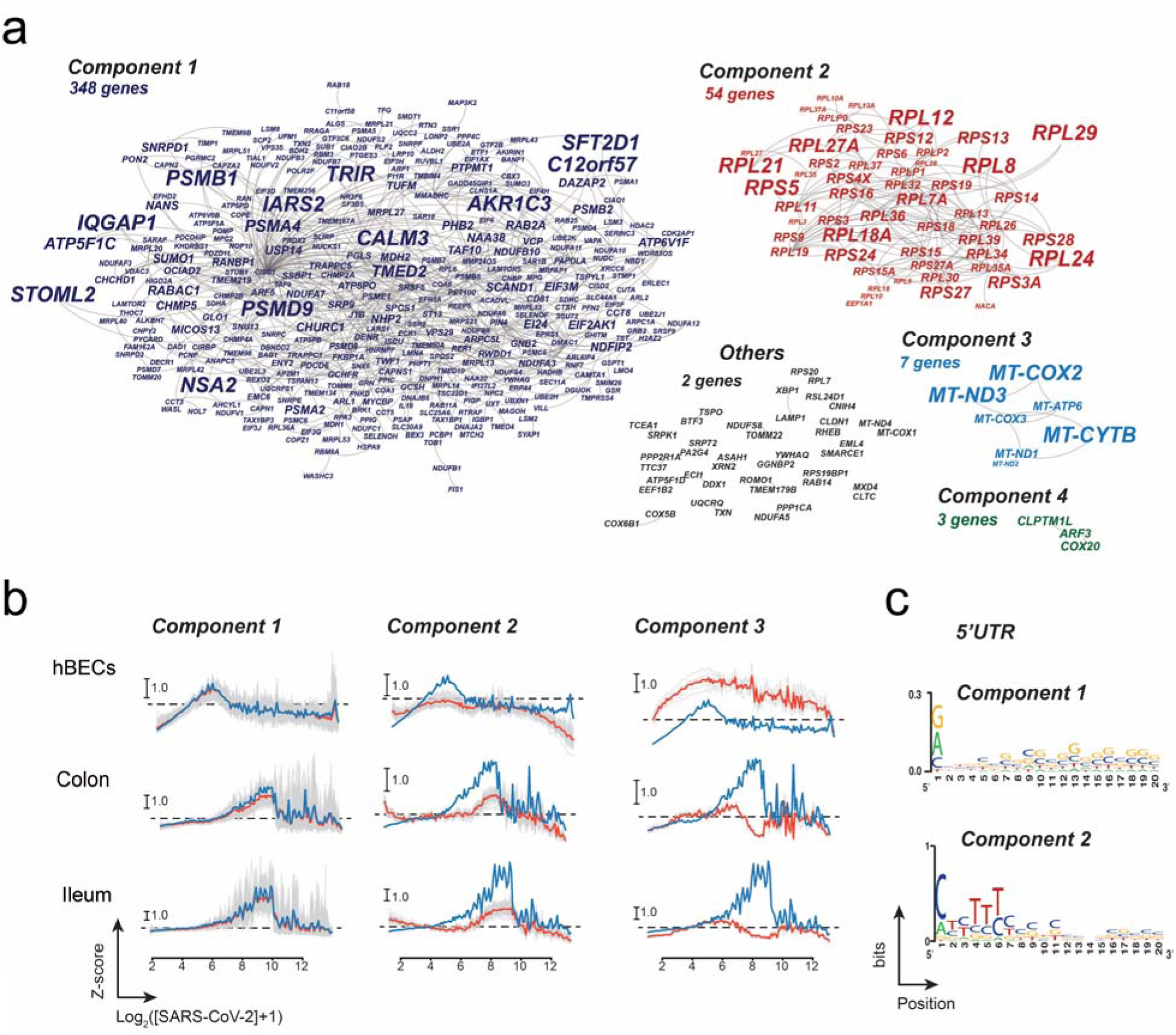
Correlation network of the common transcriptional response to SARS-CoV-2 infection. **a** Identification of correlated transcriptional patterns from the hBECS, colon and ileum datasets resulted in network comprising 25 connected components or modules. **b** Transcriptional profiles of the three largest modules. Profiles of individual genes are colored gray; the average transcriptional response of each module is shown in red. For comparison, the mean response profile is shown in blue. **c** Sequence logo of the 5’ UTRs of genes belonging to modules 1 and 2.

## Discussion

Our analysis indicates that the global transcriptional response to SARS-CoV-2 infection exhibits a behavior characterized by three phases: an initial down-regulatory phases, a middle up- regulatory phase, and a late down-regulatory phase. Clearly, this type of response cannot be analyzed using the standard two-group comparisons used in bulked RNA-seq methods (*e.g*., volcano plots) (Brennecke et al., 2013; Vallejos et al., 2016; Slovin et al., 2021). This is of special concern in the case of viral infection studies, as macroscopic timepoints (*e.g*., days or hours) comprise a complex mixture of cellular states that must be disentangled prior to running any biologically meaningful analysis. Failure to do this can introduce biases in the selection of DEGs as opposing regulatory trends in cells at different transcriptional stages might cancel-out. Analysis of the viral infection process should probably include other metrics, and as suggested here, transcriptional profiles might be a better representation of the response of individual genes along the infection timeline. Standard filtering procedures of cells in scRNA-seq are also another important source of bias, as is customary to discard cells with a low number of mapped genes and /or a high proportion of mitochondrial transcripts (Grün & van Oudenaarden, 2015; Ilicic et al., 2016; Zheng et al., 2017; Qi et al., 2020). This filtering strategy might not be adequate in studies of viral dynamics as at late infection times cells are expected to be dominated by viral transcripts and the diversity of transcribed genes is expected to be low, thus removing cells that might provide valuable information on the molecular events happening in the late stages of viral infection. In our opinion, after filtering for void-cells and multiplets, cells should be selected with respect to the number of individual transcripts and not the total number of mapped genes. A similar logic applies to the removal of cells with high levels of mitochondrial-encoded transcripts, as overexpression of mitochondrial genes might represent a legitimate response to viral infection as has been shown for SARS-CoV-2 (Singh et al., 2020; Miller et al., 2021; Yang et al., 2021).

Considering our observations, we wish to propose an updated model of SARS-CoV-2 infection. A good model must be able to explain: (*i*) the transcriptional downregulatory phase at early infection times exhibited by about ∼90% of genes, (*ii*) the selective transcriptional modulation of mitochondrial encoded genes and genes involved translation, (*iii*) the global increase in host transcript levels observed at the middle of the infection cycle, and (*iv*) the second downregulatory phase at late infection times. We believe that the multiprong strategy proposed by Finkel et al. (2021), in combination with the known biology of the viral nonstructural protein 1 (nsp1), can explain the results presented here. The multiprong strategy explains the shutdown of host protein synthesis induced by SARS-CoV-2 as a combination of three effects: a global inhibition of protein translation, degradation of cytosolic cellular transcripts and blockage of nuclear mRNA export (Finkel et al., 2021). Additionally, it is also well-stablished that nsp1 is responsible for orchestrating the transcriptional and translational shutdown induced by SARS-CoV-2. First, nsp1 is a strong inhibitor of translation affecting translation of both host and viral mRNA (Lapointe et al., 2021) by docking its C-terminal domain to the mRNA entry channel of the 40S ribosomal subunit (Schubert et al., 2020; Thoms et al., 2020; Yuan et al., 2020). Second, expression of nsp1 is sufficient to induce the global degradation of host mRNAs (Kamitami et al., 2006, 2009; Huang et al., 2011; Burke et al., 2021; Finkel et al., 2021; Lee et al., 2021) by an unknown mechanism independent of a viral encoded RNase and /or ribonuclease L (Liang et al., 2006; Burke et al., 2021). Third, immunoprecipitation and mass spectrometry studies have shown that the N-terminal domain of nsp1 interacts with the mRNA export receptor protein NFX1 preventing its binding to mRNA export adaptors that results in the accumulation of mRNA in the nucleus (Zhang et al., 2021). Clearly, nsp1 is the key factor involved in shutting down host protein expression.

The observed downregulatory phase is well-supported by many studies that have shown that soon after infection, SARS-CoV-2 mRNA transport is stalled and there is a rapid degradation of cytoplasmic mRNA (Kamitami et al., 2006, 2009; Huang et al., 2011; Burke et al., 2021; Finkel et al., 2021; Lee et al., 2021). Some believe that the inhibition of nuclear mRNA export is a direct consequence of the widespread mRNA degradation in the cytosol (Burke et al., 2021). However, we believe the opposite to be true and propose a mechanism where the down- regulatory phase results from altering the natural steady state dynamics between mRNA degradation and export to the cytoplasm (Fig. 7). The levels of mRNAs in the cytoplasm are determined by a steady-state equilibrium between the rates of transcription, nuclear export, and natural RNA turnover in the cytoplasm (Garneau et al., 2007), therefore, a global blockage of mRNA export will compromise the input of newly synthesized mRNA into the cytoplasm resulting in global decrease of cytoplasmic mRNA levels and an increase in concentration in the nucleus (Fig. 7b). In other words, expression of SARS-CoV-2 nsp1 leads to a cellular state where mRNA is stalled at the nucleus and cannot replace the mRNA being degraded in the cytoplasm by its natural turnover rate (Fig. 7b). This is supported by RNA-seq data on infected cells that revealed nuclear accumulation of mRNA and a reduction of mRNAs in the cytoplasm and increased levels of intronic reads (Banerjee et al., 2021; Finkel et al., 2021). Moreover, the mRNA export block precedes the reduction in mRNA levels and only requires expression of *nsp1* (Zhang et al., 2021); and a study of IFNB1 induction upon SARS-CoV-2 infection found that soon after infection, transcribed mRNAs fail to disseminate from transcriptional foci and are preferentially retained in the nucleus (Burke et al., 2021).

**Figure 7.**
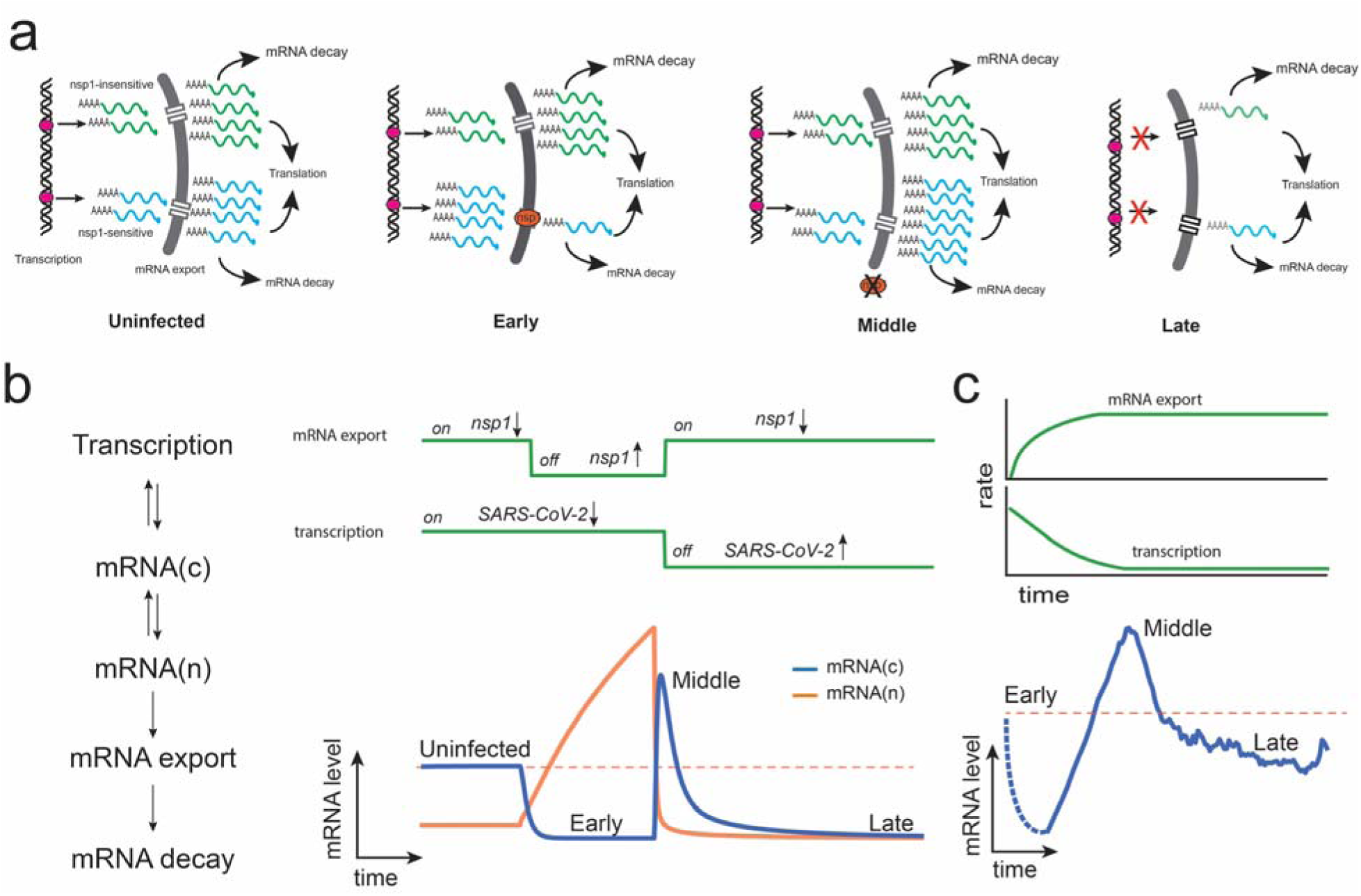
Proposed model to explain transcriptional response of infected cells to SARS-CoV-2 infection. **a** Selective nuclear export of nsp1-sensitive transcripts will result in the degradation of cytoplasmic mRNAs and their accumulation in the nucleus. Once the blockage is removed, retained transcripts are released resulting in a rapid increase of cytoplasmic mRNA levels. Shutdown of transcription during the late infection stage will result in a global decrease of host mRNAs. **b** A simplified dynamical model simulating nuclear mRNA export and transcription as on / off switches is enough to capture the transcriptional profiles observed. **c** Reinterpretation of the average transcriptional profile from hBECs. It is likely that the shutdown of transcription and nuclear export occurs gradually.

Inhibition of nuclear transport by nsp1 can also explain the differential transcriptional response of transcripts involved in translation and mitochondrial encoded transcripts. Most nuclear encoded transcripts are exported into the cytoplasm by a mechanism involving the heterodimeric export receptor NXF1·NXT1 (Xie & Ren, 2019) and it has been shown that nsp1 displaces NXF1 from the nuclear pore complex, impairing the docking of mRNA (Zhang et al. 2021). However, some transcripts can be exported by alternative pathways, indeed, regulation of mRNA export is a mechanism used modulate several critical biological processes such as DNA repair, stress response, maintenance of pluripotency (Xie & Ren, 2019). We postulate that host transcripts can be classified into nsp1-sensitive and nps1-insensitive. In our view, nsp1- sensitive transcripts correspond to transcriptionally active loci at the time of infection that are blocked by the interaction of nsp1 with NXF1. These transcripts correspond to about 90% of the transcriptome, including genes involved in the interferon response. This could explain the differential response of mRNAs involved in translation. Interestingly, a selective nuclear export mechanism of mRNAs related to translational function, including ribosomal proteins, have been shown to exist involving protein TDP-43 (Nagano et al., 2020), an RNA-binding protein involved in the regulation of transport and translation of mature mRNA in the cytoplasm (Ratti & Buratti, 2016). The differential regulation of genes involved in translation has been independently demonstrated by several studies which revealed that genes with 5’ terminal oligopyrimidine tracts tend to escape the suppression of translation induced by SARS-CoV-2 (Rao et al., 2021) and replacement of these sequences with purines results in reduced translational efficiency (Rao et al., 2021). However, we believe that possessing TOP motif is not a sufficient condition for selective transport of translational mRNAs, as several TOP containing sequences are part of the mean response subset (Supplementary Data 3). Selective mRNA export inhibition also explains the differential response of mitochondrial encoded genes as these genes do not require a nuclear export mechanism and should not be affected by the nsp1 blockage. The differential regulation of mitochondrial encoded genes is also well supported by experimental evidence that showed that they are less affected by SARS-CoV-2 infection than cellular transcripts (Miller et al., 2021; Rao et al., 2021; Finkel et al., 2021).

The up-regulatory response observed during the middle phase of SARS-CoV-2 infection is a novel and unexpected observation that deserves additional discussion. If we assume that nsp1 is degraded or inactivated after it has accomplished its function, then the stalled mRNA transcripts can be finally exported to the nucleus in a rapid burst which would look like an up- regulatory event during the intermediate infection stage (Fig. 7). This rapid burst of newly synthesized transcripts can also explain the late but strong response of the interferon system typical of COVID-19 disease (Lee et al., 2020; Wilk et al., 2020; Zhou et al., 2020). It is also reasonable to assume that, when the nuclear blockage stops, it is already too late for the host cell to outcompete the replicating virus which results in the final down-regulatory phase observed at high SARS-CoV-2 levels. Qualitatively, this behavior can be simulated with a simple dynamical model where global mRNA export and transcription act like switches (Fig. 7a). Expression of *nsp1* acts like an off-switch for nsp1-sensitive transcripts that results in their accumulation in the nucleus while nsp1-insensitive transcripts can transit normally to the cytoplasm. The middle phase corresponds to the inactivation of the nsp1-induced blockage which releases the stalled transcript to the cytoplasm that results in the increase transcription levels observed during the middle phase. Finally, the late phase corresponds to a shutdown of transcription than also results in a global decrease in transcript levels.

In summary, our analysis reveals a complex transcriptional response to SARS-CoV-2 infection that can be easily explained using the blockage of mRNA transport as the key molecular event. We believe that this model provides a parsimonious explanation of the host transcriptional and translational shutdown induced by SARS-CoV-2 and suggests that targeting the nsp1 ability to disrupt nuclear export might be the key to counteract the effect of this virus within infected cells.

## Acknowledgements

We thank Sergi Valverde, Salvador Durán-Nebreda, the members of the I^2^SysBio-CRM UA DisCoVir, and our labmates for useful suggestions and productive discussions. This work was supported by European Commission – NextGenerationEU (Regulation EU 2020 /2094) through CSIC’s Global Health Platform (PTI+ Salud Global) grants SGL2021-03-009 and SGL2021-03-052 to S.F.E.

## Supplementary information

**Supplementary Figure 1.**
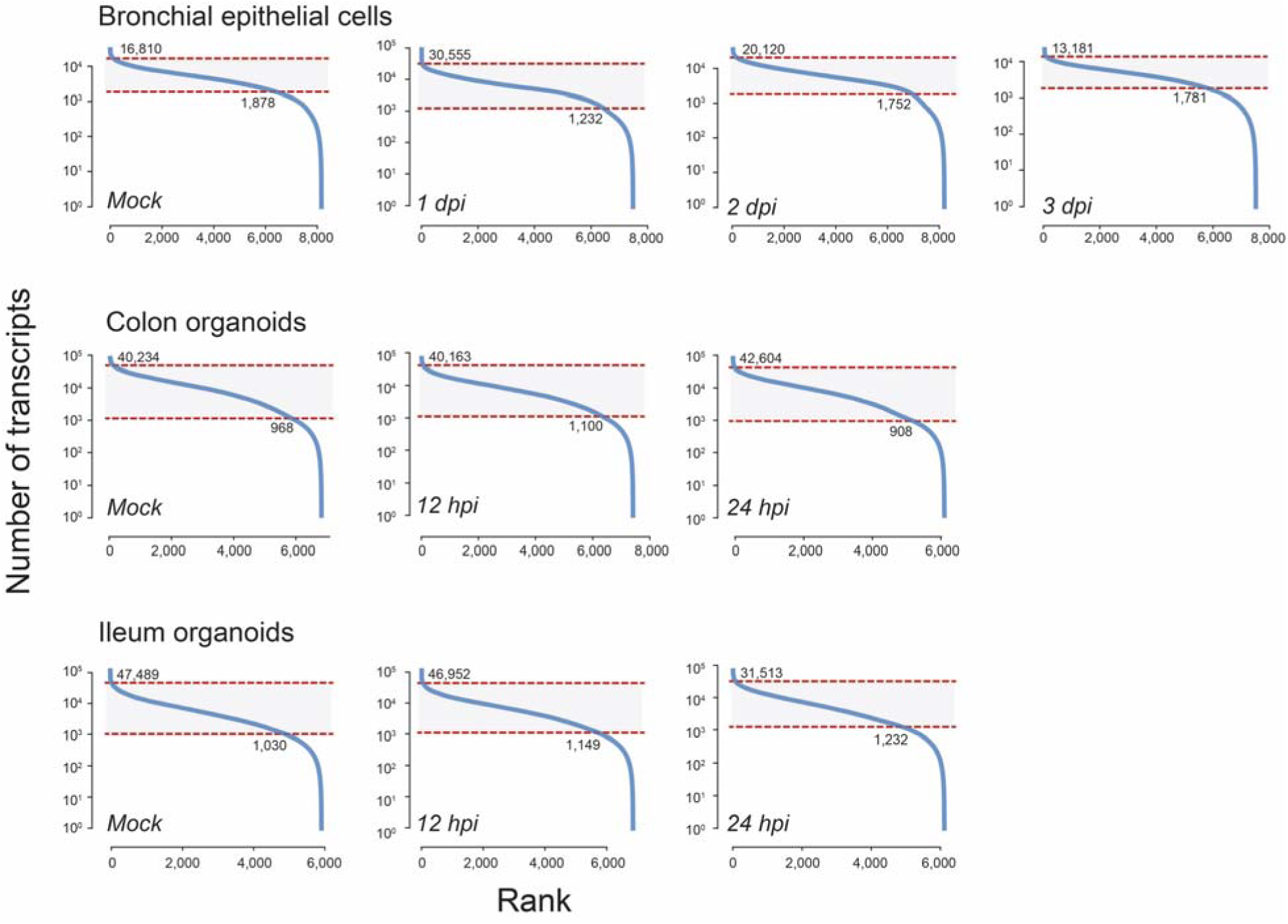
Filtering of void GEMs and multiplets. GEMs were ordered with respect to the number of transcripts the local standard deviation of the log_10_ using window size of five datapoints. Upper and lower thresholds delimiting multiples and voids GEMS were determined using 1.5×IQR rule. The cells used in this study correspond to the grey area.

**Supplementary Figure 2.**
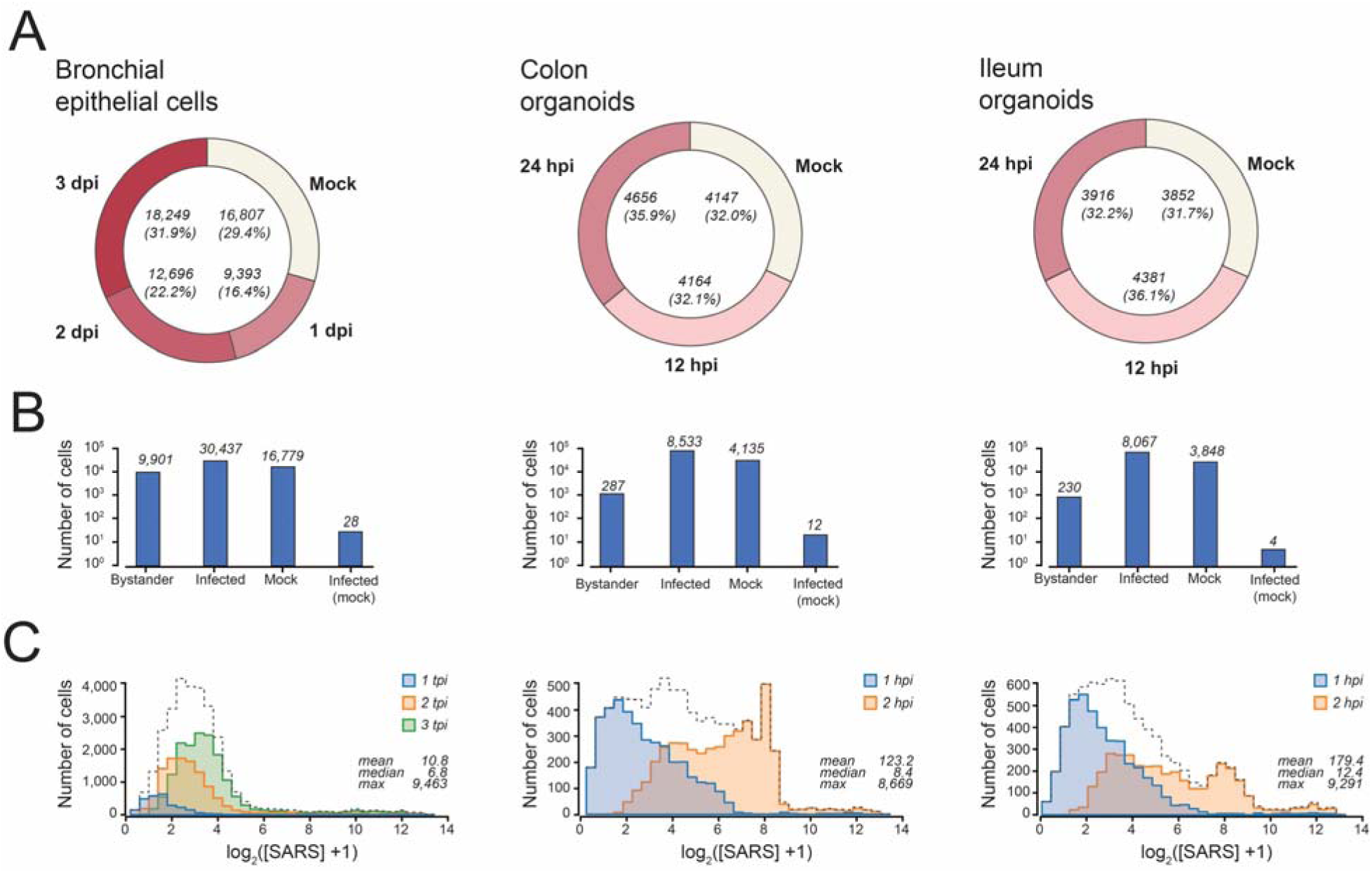
Global composition of the scRNA-seq datasets. **a** Proportion of GEMs from mock and infected treatments from each dataset. **b** Proportion of bystanders, infected, mock, and infected mock cells in each dataset. **c** Distribution of cells with respect to viral loads for each treatment. Statistics presented in these plots correspond to observed counts.

**Supplementary Table 1.**
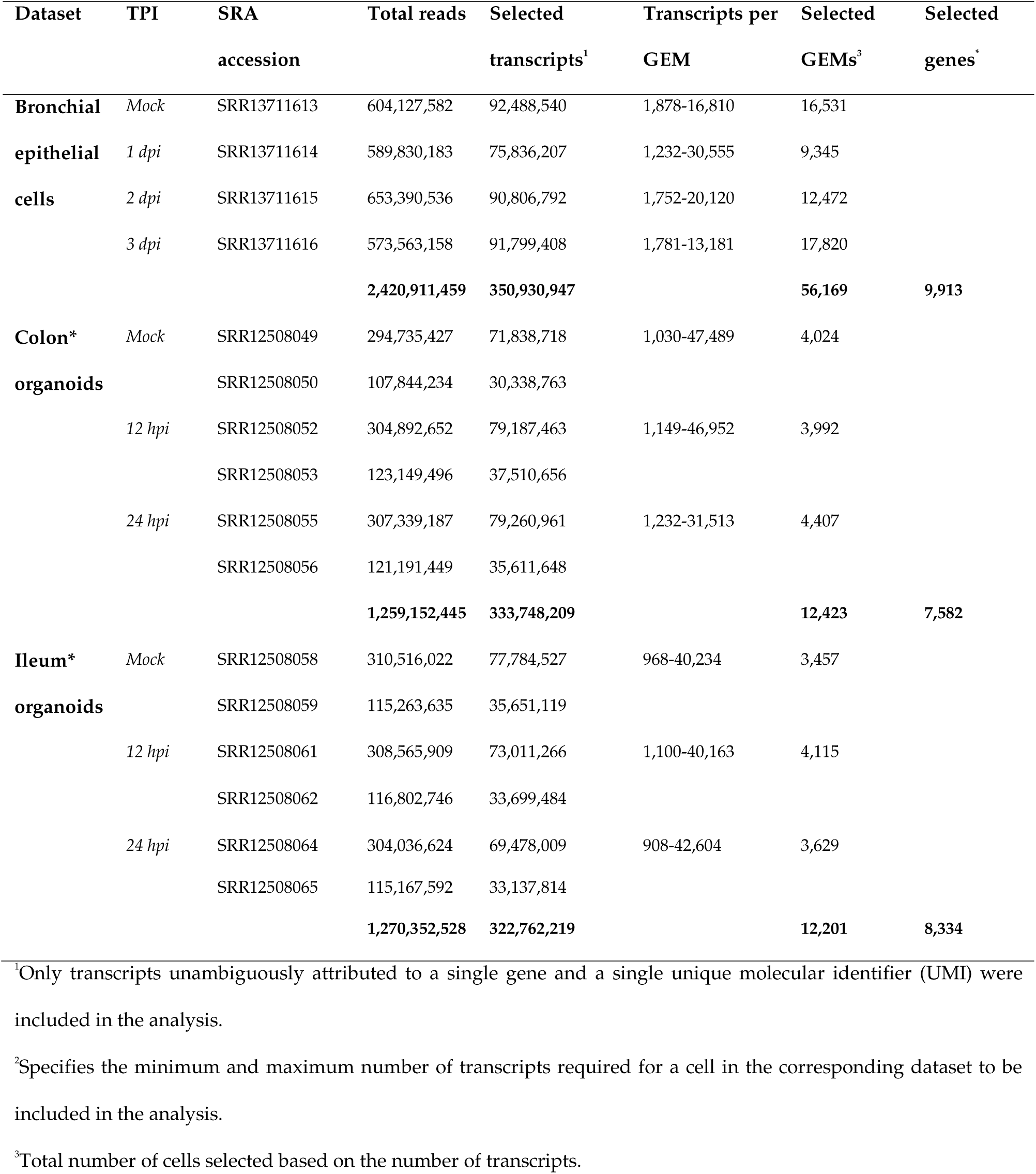
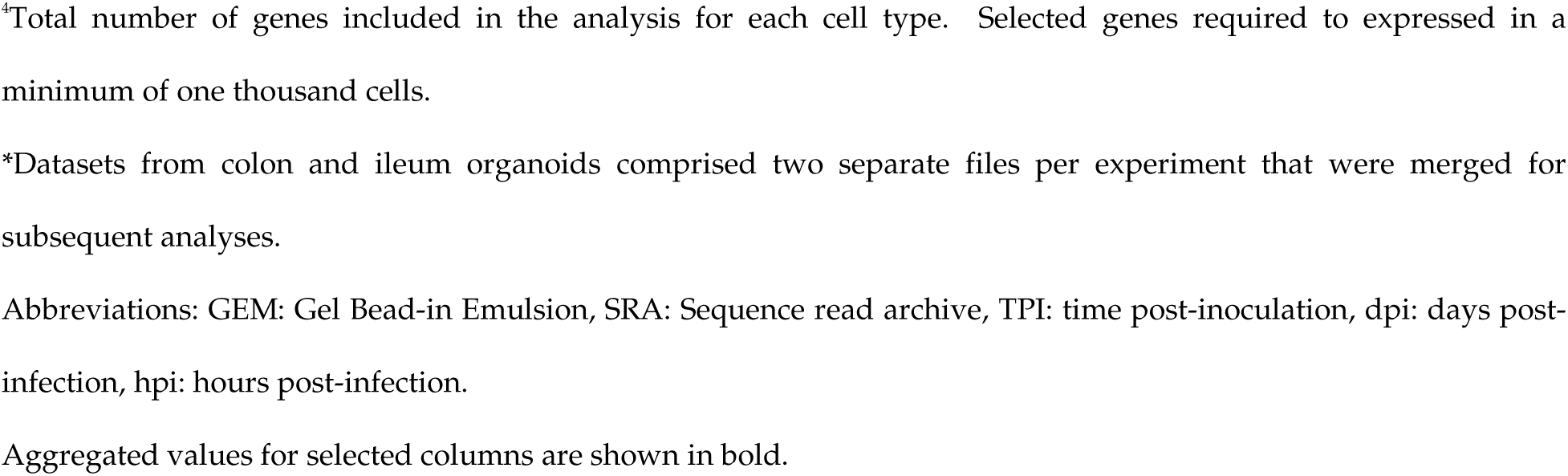
Global statistics of the dataset used in the analysis.

| Dataset | TPI | SRA<br>accession | Total reads | Selected<br>transcripts<sup>1</sup> | Transcripts per<br>GEM | Selected<br>GEMs<sup>3</sup> | Selected<br>genes<sup>2</sup> |
| --- | --- | --- | --- | --- | --- | --- | --- |
| **Bronchial** | *Mock* | SRR13711613 | 604,127,582 | 92,488,540 | 1,878-16,810 | 16,531 |  |
| **epithelial** | *1 dpi* | SRR13711614 | 589,830,183 | 75,836,207 | 1,232-30,555 | 9,345 |  |
| **cells** | *2 dpi* | SRR13711615 | 653,390,536 | 90,806,792 | 1,752-20,120 | 12,472 |  |
|  | *3 dpi* | SRR13711616 | 573,563,158 | 91,799,408 | 1,781-13,181 | 17,820 |  |
|  |  |  | **2,420,911,459** | **350,930,947** |  | **56,169** | **9,913** |
| **Colon\*** | *Mock* | SRR12508049 | 294,735,427 | 71,838,718 | 1,030-47,489 | 4,024 |  |
| **organoids** |  | SRR12508050 | 107,844,234 | 30,338,763 |  |  |  |
|  | *12 hpi* | SRR12508052 | 304,892,652 | 79,187,463 | 1,149-46,952 | 3,992 |  |
|  |  | SRR12508053 | 123,149,496 | 37,510,656 |  |  |  |
|  | *24 hpi* | SRR12508055 | 307,339,187 | 79,260,961 | 1,232-31,513 | 4,407 |  |
|  |  | SRR12508056 | 121,191,449 | 35,611,648 |  |  |  |
|  |  |  | **1,259,152,445** | **333,748,209** |  | **12,423** | **7,582** |
| **Ileum\*** | *Mock* | SRR12508058 | 310,516,022 | 77,784,527 | 968-40,234 | 3,457 |  |
| **organoids** |  | SRR12508059 | 115,263,635 | 35,651,119 |  |  |  |
|  | *12 hpi* | SRR12508061 | 308,565,909 | 73,011,266 | 1,100-40,163 | 4,115 |  |
|  |  | SRR12508062 | 116,802,746 | 33,699,484 |  |  |  |
|  | *24 hpi* | SRR12508064 | 304,036,624 | 69,478,009 | 908-42,604 | 3,629 |  |
|  |  | SRR12508065 | 115,167,592 | 33,137,814 |  |  |  |
|  |  |  | **1,270,352,528** | **322,762,219** |  | **12,201** | **8,334** |

<sup>1</sup>Only transcripts unambiguously attributed to a single gene and a single unique molecular identifier (UMI) were included in the analysis.
<sup>2</sup>Specifies the minimum and maximum number of transcripts required for a cell in the corresponding dataset to be included in the analysis.
<sup>3</sup>Total number of cells selected based on the number of transcripts.
<sup>a</sup>Total number of genes included in the analysis for each cell type. Selected genes required to expressed in a minimum of one thousand cells.
\*Datasets from colon and ileum organoids comprised two separate files per experiment that were merged for subsequent analyses.
Abbreviations: GEM: Gel Bead-in Emulsion, SRA: Sequence read archive, TPI: time post-inoculation, dpi: days post-infection, hpi: hours post-infection.
Aggregated values for selected columns are shown in bold.

**Supplementary Data 1.** Database of reference human messenger RNAs and SARS-CoV-2 genomes used for mapping reads. Raw count matrices.

**Supplementary Data 2.** Expression matrices and transcript levels in uninfected cells.

**Supplementary Data 3.** Differential Expressed Gene analysis used in the volcano plots analysis. Datasets and results used in the Gene Ontology analyses. Classification of outliers.

**Supplementary Data 4.** Transcriptional profiles for all genes in each cell type.

**Supplementary Data 5.** Networks used in the analyses presented in figures 5 and 6.

## References

Addetia, A., Lieberman, N., Phung, Q., Hsiang, T. Y., Xie, H., Roychoudhury, P., Shrestha, L., Loprieno, M. A., Huang, M. L., Gale, M., Jr, Jerome, K. R., & Greninger, A. L. (2021). SARS-CoV-2 ORF6 disrupts bidirectional nucleocytoplasmic transport through interactions with rae1 and nup98. mBio, 12(2), e00065–21. https://doi.org/10.1128/mBio.00065-21

Aric A. Hagberg, Daniel A. Schult and Pieter J. Swart, “Exploring network structure, dynamics, and function using NetworkX”, in Proceedings of the 7th Python in Science Conference (SciPy2008), Gäel Varoquaux, Travis Vaught, and Jarrod Millman (Eds), (Pasadena, CA USA), pp. 11–15, Aug 2008

Banerjee, A. K., Blanco, M. R., Bruce, E. A., Honson, D. D., Chen, L. M., Chow, A., Bhat, P., Ollikainen, N., Quinodoz, S. A., Loney, C., Thai, J., Miller, Z. D., Lin, A. E., Schmidt, M. M., Stewart, D. G., Goldfarb, D., De Lorenzo, G., Rihn, S. J., Voorhees, R. M., Botten, J. W., … Guttman, M. (2020). SARS-CoV-2 disrupts splicing, translation, and protein trafficking to suppress host defenses. Cell, 183(5), 1325–1339.e21. https://doi.org/10.1016/j.cell.2020.10.004

Barrantes F. J. (2021). The contribution of biophysics and structural biology to current advances in COVID-19. Annual Review of Biophysics, 50, 493–523. https://doi.org/10.1146/annurev-biophys-102620-080956

Bastian, M., Heymann, S., & Jacomy, M. (2009). Gephi: an open source software for exploring and manipulating networks. In Third international AAAI conference on weblogs and social media.

Benjamini, Y. and Hochberg, Y. (1995). Controlling the false discovery rate: A practical and powerful approach to multiple testing. Journal of the Royal Statistical Society: Series B (Methodological), 57: 289–300. https://doi.org/10.1111/j.2517-6161.1995.tb02031.x

Boratyn, G. M., Thierry-Mieg, J., Thierry-Mieg, D., Busby, B., & Madden, T. L. (2019). Magic-BLAST, an accurate RNA-seq aligner for long and short reads. BMC Bioinformatics, 20(1), 405. https://doi.org/10.1186/s12859-019-2996-x

Brant, A. C., Tian, W., Majerciak, V., Yang, W., & Zheng, Z. M. (2021). SARS-CoV-2: from its discovery to genome structure, transcription, and replication. Cell & Bioscience, 11(1), 136. https://doi.org/10.1186/s13578-021-00643-z

Brennecke, P., Anders, S., Kim, J. K., Kołodziejczyk, A. A., Zhang, X., Proserpio, V., Baying, B., Benes, V., Teichmann, S. A., Marioni, J. C., & Heisler, M. G. (2013). Accounting for technical noise in single-cell RNA-seq experiments. Nature Methods, 10(11), 1093–1095. https://doi.org/10.1038/nmeth.2645

Burke, J. M., St Clair, L. A., Perera, R., & Parker, R. (2021). SARS-CoV-2 infection triggers widespread host mRNA decay leading to an mRNA export block. RNA, 27(11), 1318–1329. https://doi.org/10.1261/rna.078923.121

Chen, G., Ning, B., & Shi, T. (2019). Single-Cell RNA-Seq technologies and related computational data analysis. Frontiers in Genetics, 10, 317. https://doi.org/10.3389/fgene.2019.00317

Ciuffi, A., Rato, S., & Telenti, A. (2016). Single-cell genomics for virology. Viruses, 8(5), 123. https://doi.org/10.3390/v8050123

Clayton, S. A., Daley, K. K., MacDonald, L., Fernandez-Vizarra, E., Bottegoni, G., O’Neil, J. D., Major, T., Griffin, D., Zhuang, Q., Adewoye, A. B., Woolcock, K., Jones, S. W., Goodyear, C., Elmesmari, A., Filer, A., Tennant, D. A., Alivernini, S., Buckley, C. D., Pitceathly, R., Kurowska-Stolarska, M., … Clark, A. R. (2021). Inflammation causes remodeling of mitochondrial cytochrome c oxidase mediated by the bifunctional gene *C15orf48*. Science Advances, 7(50), eabl5182. https://doi.org/10.1126/sciadv.abl5182

Cristinelli, S., & Ciuffi, A. (2018). The use of single-cell RNA-Seq to understand virus-host interactions. Current Opinion in Virology, 29, 39–50. https://doi.org/10.1016/j.coviro.2018.03.001

Ding, J., Adiconis, X., Simmons, S. K., Kowalczyk, M. S., Hession, C. C., Marjanovic, N. D., Hughes, T. K., Wadsworth, M. H., Burks, T., Nguyen, L. T., Kwon, J., Barak, B., Ge, W., Kedaigle, A. J., Carroll, S., Li, S., Hacohen, N., Rozenblatt-Rosen, O., Shalek, A. K., Villani, A. C., … Levin, J. Z. (2020). Systematic comparison of single-cell and single-nucleus RNA-sequencing methods. Nature Biotechnology, 38(6), 737–746. https://doi.org/10.1038/s41587-020-0465-8

Ding, J., Sharon, N., & Bar-Joseph, Z. (2022). Temporal modelling using single-cell transcriptomics. Nature Reviews Genetics, 23(6), 355–368. https://doi.org/10.1038/s41576-021-00444-7

Finkel, Y., Gluck, A., Nachshon, A., Winkler, R., Fisher, T., Rozman, B., Mizrahi, O., Lubelsky, Y., Zuckerman, B., Slobodin, B., Yahalom-Ronen, Y., Tamir, H., Ulitsky, I., Israely, T., Paran, N., Schwartz, M., & Stern-Ginossar, N. (2021). SARS-CoV-2 uses a multipronged strategy to impede host protein synthesis. Nature, 594(7862), 240–245. https://doi.org/10.1038/s41586-021-03610-3

Garneau, N. L., Wilusz, J., & Wilusz, C. J. (2007). The highways and byways of mRNA decay. Nature Reviews Molecular Cell Biology, 8(2), 113–126. https://doi.org/10.1038/nrm2104

Geller, R., Taguwa, S., & Frydman, J. (2012). Broad action of Hsp90 as a host chaperone required for viral replication. Biochimica et Biophysica Acta, 1823(3), 698–706. https://doi.org/10.1016/j.bbamcr.2011.11.007

Griffiths, J. A., Scialdone, A., & Marioni, J. C. (2018). Using single-cell genomics to understand developmental processes and cell fate decisions. Molecular Systems Biology, 14(4), e8046. https://doi.org/10.15252/msb.20178046

Grün, D., & van Oudenaarden, A. (2015). Design and analysis of single-cell sequencing experiments. Cell, 163(4), 799–810. https://doi.org/10.1016/j.cell.2015.10.039

Hu, B., Guo, H., Zhou, P., & Shi, Z. L. (2021). Characteristics of SARS-CoV-2 and COVID-19. Nature Reviews Microbiology, 19(3), 141–154. https://doi.org/10.1038/s41579-020-00459-7

Huang, C., Lokugamage, K. G., Rozovics, J. M., Narayanan, K., Semler, B. L., & Makino, S. (2011). SARS coronavirus nsp1 protein induces template-dependent endonucleolytic cleavage of mRNAs: viral mRNAs are resistant to nsp1-induced RNA cleavage. PLoS Pathogens, 7(12), e1002433. https://doi.org/10.1371/journal.ppat.1002433

Hwang, B., Lee, J. H., & Bang, D. (2018). Single-cell RNA sequencing technologies and bioinformatics pipelines. Experimental & Molecular Medicine, 50(8), 1–14. https://doi.org/10.1038/s12276-018-0071-8

Ilicic, T., Kim, J. K., Kolodziejczyk, A. A., Bagger, F. O., McCarthy, D. J., Marioni, J. C., & Teichmann, S. A. (2016). Classification of low quality cells from single-cell RNA-seq data. Genome Biology, 17, 29. https://doi.org/10.1186/s13059-016-0888-1

Islam, S., Zeisel, A., Joost, S., La Manno, G., Zajac, P., Kasper, M., Lönnerberg, P., & Linnarsson, S. (2014). Quantitative single-cell RNA-seq with unique molecular identifiers. Nature Methods, 11(2), 163–166. https://doi.org/10.1038/nmeth.2772

Jin, W., Jin, W., & Pan, D. (2018). Ifi27 is indispensable for mitochondrial function and browning in adipocytes. Biochemical and Biophysical Research Communications, 501(1), 273–279. https://doi.org/10.1016/j.bbrc.2018.04.234

Kamitani, W., Narayanan, K., Huang, C., Lokugamage, K., Ikegami, T., Ito, N., Kubo, H., & Makino, S. (2006). Severe acute respiratory syndrome coronavirus nsp1 protein suppresses host gene expression by promoting host mRNA degradation. Proceedings of the National Academy of Sciences of the United States of America, 103(34), 12885–12890. https://doi.org/10.1073/pnas.0603144103

Kamitani, W., Huang, C., Narayanan, K., Lokugamage, K. G., & Makino, S. (2009). A two-pronged strategy to suppress host protein synthesis by SARS coronavirus Nsp1 protein. Nature Structural & Molecular Biology, 16(11), 1134–1140. https://doi.org/10.1038/nsmb.1680

Kharchenko P. V. (2021). The triumphs and limitations of computational methods for scRNA-seq. Nature Methods, 18(7), 723–732. https://doi.org/10.1038/s41592-021-01171-x

Kumar, A., Ishida, R., Strilets, T., Cole, J., Lopez-Orozco, J., Fayad, N., Felix-Lopez, A., Elaish, M., Evseev, D., Magor, K. E., Mahal, L. K., Nagata, L. P., Evans, D. H., & Hobman, T. C. (2021). SARS-CoV-2 nonstructural protein 1 inhibits the interferon response by causing depletion of key host signaling factors. Journal of Virology, 95(13), e0026621. https://doi.org/10.1128/JVI.00266-21

Lambert, S. A., Jolma, A., Campitelli, L. F., Das, P. K., Yin, Y., Albu, M., Chen, X., Taipale, J., Hughes, T. R., & Weirauch, M. T. (2018). The human transcription factors. Cell, 172(4), 650– 665. https://doi.org/10.1016/j.cell.2018.01.029

Lapointe, C. P., Grosely, R., Johnson, A. G., Wang, J., Fernández, I. S., & Puglisi, J. D. (2021). Dynamic competition between SARS-CoV-2 NSP1 and mRNA on the human ribosome inhibits translation initiation. Proceedings of the National Academy of Sciences of the United States of America, 118(6), e2017715118. https://doi.org/10.1073/pnas.2017715118

Lee, W. S., Yousefi, M., Yan, B., Yong, C. L., & Ooi, Y. S. (2021). Know your enemy and know yourself - the case of SARS-CoV-2 host factors. Current Opinion in Virology, 50, 159–170. https://doi.org/10.1016/j.coviro.2021.08.007

Liang, S. L., Quirk, D., & Zhou, A. (2006). RNase L: its biological roles and regulation. IUBMB Life, 58(9), 508–514. https://doi.org/10.1080/15216540600838232

Liao, M., Liu, Y., Yuan, J., Wen, Y., Xu, G., Zhao, J., Cheng, L., Li, J., Wang, X., Wang, F., Liu, L., Amit, I., Zhang, S., & Zhang, Z. (2020). Single-cell landscape of bronchoalveolar immune cells in patients with COVID-19. Nature Medicine, 26(6), 842–844. https://doi.org/10.1038/s41591-020-0901-9

Liao, J. Y., Yang, B., Zhang, Y. C., Wang, X. J., Ye, Y., Peng, J. W., Yang, Z. Z., He, J. H., Zhang, Y., Hu, K., Lin, D. C., & Yin, D. (2020). EuRBPDB: a comprehensive resource for annotation, functional and oncological investigation of eukaryotic RNA binding proteins (RBPs). Nucleic Acids Research, 48(D1), D307–D313. https://doi.org/10.1093/nar/gkz823

Liu, X. Y., Chen, W., Wei, B., Shan, Y. F., & Wang, C. (2011). IFN-induced TPR protein IFIT3 potentiates antiviral signaling by bridging MAVS and TBK1. Journal of Immunology, 187(5), 2559–2568. https://doi.org/10.4049/jimmunol.1100963

Lyabin, D. N., Eliseeva, I. A., & Ovchinnikov, L. P. (2014). YB-1 protein: functions and regulation. RNA, 5(1), 95–110. https://doi.org/10.1002/wrna.1200

Martin-Sancho, L., Lewinski, M. K., Pache, L., Stoneham, C. A., Yin, X., Becker, M. E., Pratt, D., Churas, C., Rosenthal, S. B., Liu, S., Weston, S., De Jesus, P. D., O’Neill, A. M., Gounder, A. P., Nguyen, C., Pu, Y., Curry, H. M., Oom, A. L., Miorin, L., Rodriguez-Frandsen, A., … Chanda, S. K. (2021). Functional landscape of SARS-CoV-2 cellular restriction. Molecular Cell, 81(12), 2656–2668.e8. https://doi.org/10.1016/j.molcel.2021.04.008

McWilliam Leitch, E. C., & McLauchlan, J. (2013). Determining the cellular diversity of hepatitis C virus quasispecies by single-cell viral sequencing. Journal of Virology, 87(23), 12648–12655. https://doi.org/10.1128/JVI.01602-13

Meng, Q., & Xia, Y. (2011). c-Jun, at the crossroad of the signaling network. Protein & Cell, 2(11), 889–898. https://doi.org/10.1007/s13238-011-1113-3

Meyuhas, O., & Kahan, T. (2015). The race to decipher the top secrets of TOP mRNAs. Biochimica et Biophysica Acta, 1849(7), 801–811. https://doi.org/10.1016/j.bbagrm.2014.08.015

Miller, B., Silverstein, A., Flores, M., Cao, K., Kumagai, H., Mehta, H. H., Yen, K., Kim, S. J., & Cohen, P. (2021). Host mitochondrial transcriptome response to SARS-CoV-2 in multiple cell models and clinical samples. Scientific Reports, 11(1), 3. https://doi.org/10.1038/s41598-020-79552-z

Miorin, L., Kehrer, T., Sanchez-Aparicio, M. T., Zhang, K., Cohen, P., Patel, R. S., Cupic, A., Makio, T., Mei, M., Moreno, E., Danziger, O., White, K. M., Rathnasinghe, R., Uccellini, M., Gao, S., Aydillo, T., Mena, I., Yin, X., Martin-Sancho, L., Krogan, N. J., … García-Sastre, A. (2020). SARS-CoV-2 ORF6 hijacks nup98 to block STAT nuclear import and antagonize interferon signaling. Proceedings of the National Academy of Sciences of the United States of America, 117(45), 28344–28354. https://doi.org/10.1073/pnas.2016650117

Miyake, T., & McDermott, J. C. (2022). Nucleolar localization of c-Jun. The FEBS Journal, 289(3), 748–765. https://doi.org/10.1111/febs.16187

Nagano, S., Jinno, J., Abdelhamid, R. F., Jin, Y., Shibata, M., Watanabe, S., Hirokawa, S., Nishizawa, M., Sakimura, K., Onodera, O., Okada, H., Okada, T., Saito, Y., Takahashi-Fujigasaki, J., Murayama, S., Wakatsuki, S., Mochizuki, H., & Araki, T. (2020). TDP-43 transports ribosomal protein mRNA to regulate axonal local translation in neuronal axons. Acta Neuropathologica, 140(5), 695–713. https://doi.org/10.1007/s00401-020-02205-y

Nagasu, S., Sudo, T., Kinugasa, T., Yomoda, T., Fujiyoshi, K., Shigaki, T., & Akagi, Y. (2019). Y•box•binding protein 1 inhibits apoptosis and upregulates EGFR in colon cancer. Oncology Reports, 41(5), 2889–2896. https://doi.org/10.3892/or.2019.7038

Nakagawa, K., Narayanan, K., Wada, M., Popov, V. L., Cajimat, M., Baric, R. S., & Makino, S. (2018). The endonucleolytic RNA cleavage function of nsp1 of Middle East respiratory syndrome coronavirus promotes the production of infectious virus particles in specific human cell lines. Journal of Virology, 92(21), e01157–18. https://doi.org/10.1128/JVI.01157-18

Qi, R., Ma, A., Ma, Q., & Zou, Q. (2020). Clustering and classification methods for single-cell RNA-sequencing data. Briefings in Bioinformatics, 21(4), 1196–1208. https://doi.org/10.1093/bib/bbz062

Ramirez Alvarez, C., Kee, C., Sharma, A. K., Thomas, L., Schmidt, F. I., Stanifer, M. L., Boulant, S., & Herrmann, C. (2021). The endogenous cellular protease inhibitor SPINT2 controls SARS-CoV-2 viral infection and is associated to disease severity. PLoS Pathogens, 17(6), e1009687. https://doi.org/10.1371/journal.ppat.1009687

Rao, S., Hoskins, I., Tonn, T., Garcia, P. D., Ozadam, H., Sarinay Cenik, E., & Cenik, C. (2021). Genes with 5’ terminal oligopyrimidine tracts preferentially escape global suppression of translation by the SARS-CoV-2 Nsp1 protein. RNA 27(9), 1025–1045. https://doi.org/10.1261/rna.078661.120

Ratti, A., & Buratti, E. (2016). Physiological functions and pathobiology of TDP-43 and FUS/TLS proteins. Journal of Neurochemistry, 138 Suppl 1, 95–111. https://doi.org/10.1111/jnc.13625

Ravindra, N. G., Alfajaro, M. M., Gasque, V., Huston, N. C., Wan, H., Szigeti-Buck, K., Yasumoto, Y., Greaney, A. M., Habet, V., Chow, R. D., Chen, J. S., Wei, J., Filler, R. B., Wang, B., Wang, G., Niklason, L. E., Montgomery, R. R., Eisenbarth, S. C., Chen, S., Williams, A., … Wilen, C. B. (2021). Single-cell longitudinal analysis of SARS-CoV-2 infection in human airway epithelium identifies target cells, alterations in gene expression, and cell state changes. PLoS Biology, 19(3), e3001143. https://doi.org/10.1371/journal.pbio.3001143

Rebendenne, A., Valadão, A., Tauziet, M., Maarifi, G., Bonaventure, B., McKellar, J., Planès, R., Nisole, S., Arnaud-Arnould, M., Moncorgé, O., & Goujon, C. (2021). SARS-CoV-2 triggers an MDA-5-dependent interferon response which is unable to control replication in lung epithelial cells. Journal of Virology, 95(8), e02415–20. https://doi.org/10.1128/JVI.02415-20

Rostom, R., Svensson, V., Teichmann, S. A., & Kar, G. (2017). Computational approaches for interpreting scRNA-seq data. FEBS Letters, 591(15), 2213–2225. https://doi.org/10.1002/1873-3468.12684

Ruiz, E. J., Lan, L., Diefenbacher, M. E., Riising, E. M., Da Costa, C., Chakraborty, A., Hoeck, J. D., Spencer-Dene, B., Kelly, G., David, J. P., Nye, E., Downward, J., & Behrens, A. (2021). JunD, not c-Jun, is the AP-1 transcription factor required for ras-induced lung cancer. JCI Insight, 6(13), e124985. https://doi.org/10.1172/jci.insight.124985

Russell, A. B., Trapnell, C., & Bloom, J. D. (2018). Extreme heterogeneity of influenza virus infection in single cells. eLife, 7, e32303. https://doi.org/10.7554/eLife.32303

Sampaio, N. G., Chauveau, L., Hertzog, J., Bridgeman, A., Fowler, G., Moonen, J. P., Dupont, M., Russell, R. A., Noerenberg, M., & Rehwinkel, J. (2021). The RNA sensor MDA5 detects SARS-CoV-2 infection. Scientific Reports, 11(1), 13638. https://doi.org/10.1038/s41598-021-92940-3

Shnayder, M., Nachshon, A., Krishna, B., Poole, E., Boshkov, A., Binyamin, A., Maza, I., Sinclair, J., Schwartz, M., & Stern-Ginossar, N. (2018). Defining the transcriptional landscape during cytomegalovirus latency with single-cell RNA sequencing. mBio, 9(2), e00013–18. https://doi.org/10.1128/mBio.00013-18

Schoggins, J. W., Wilson, S. J., Panis, M., Murphy, M. Y., Jones, C. T., Bieniasz, P., & Rice, C. M. (2011). A diverse range of gene products are effectors of the type I interferon antiviral response. Nature, 472(7344), 481–485. https://doi.org/10.1038/nature09907

Schubert, K., Karousis, E. D., Jomaa, A., Scaiola, A., Echeverria, B., Gurzeler, L. A., Leibundgut, M., Thiel, V., Mühlemann, O., & Ban, N. (2020). SARS-CoV-2 Nsp1 binds the ribosomal mRNA channel to inhibit translation. Nature Structural & Molecular Biology, 27(10), 959– 966. https://doi.org/10.1038/s41594-020-0511-8

Schuler, B. A., Habermann, A. C., Plosa, E. J., Taylor, C. J., Jetter, C., Negretti, N. M., Kapp, M. E., Benjamin, J. T., Gulleman, P., Nichols, D. S., Braunstein, L. Z., Hackett, A., Koval, M., Guttentag, S. H., Blackwell, T. S., Webber, S. A., Banovich, N. E., Vanderbilt COVID-19 Consortium Cohort, Human Cell Atlas Biological Network, Kropski, J. A., … Sucre, J. M. (2021). Age-determined expression of priming protease TMPRSS2 and localization of SARS-CoV-2 in lung epithelium. The Journal of Clinical Investigation, 131(1), e140766. https://doi.org/10.1172/JCI140766

Sen, K., Datta, S., Ghosh, A., Jha, A., Ahad, A., Chatterjee, S., Suranjika, S., Sengupta, S., Bhattacharya, G., Shriwas, O., Avula, K., Kshatri, J., Prasad, P., Swain, R., Parida, A. K., & Raghav, S. K. (2021). Single-cell immunogenomic approach identified SARS-CoV-2 protective immune signatures in asymptomatic direct contacts of COVID-19 cases. Frontiers in Immunology, 12, 733539. https://doi.org/10.3389/fimmu.2021.733539

Simeoni, M., Cavinato, T., Rodriguez, D., & Gatfield, D. (2021). I(nsp1)ecting SARS-CoV-2-ribosome interactions. Communications Biology, 4(1), 715. https://doi.org/10.1038/s42003-021-02265-0

Singh, K. K., Chaubey, G., Chen, J. Y., & Suravajhala, P. (2020). Decoding SARS-CoV-2 hijacking of host mitochondria in COVID-19 pathogenesis. American Journal of Physiology. Cell physiology, 319(2), C258–C267. https://doi.org/10.1152/ajpcell.00224.2020

Slovin, S., Carissimo, A., Panariello, F., Grimaldi, A., Bouché, V., Gambardella, G., & Cacchiarelli, D. (2021). Single-cell RNA sequencing analysis: A step-by-step overview. Methods in Molecular Biology, 2284, 343–365. https://doi.org/10.1007/978-1-0716-1307-8_19

Thoms, M., Buschauer, R., Ameismeier, M., Koepke, L., Denk, T., Hirschenberger, M., Kratzat, H., Hayn, M., Mackens-Kiani, T., Cheng, J., Straub, J. H., Stürzel, C. M., Fröhlich, T., Berninghausen, O., Becker, T., Kirchhoff, F., Sparrer, K., & Beckmann, R. (2020). Structural basis for translational shutdown and immune evasion by the Nsp1 protein of SARS-CoV-2. Science, 369(6508), 1249–1255. https://doi.org/10.1126/science.abc8665

Thorne, L. G., Reuschl, A. K., Zuliani-Alvarez, L., Whelan, M., Turner, J., Noursadeghi, M., Jolly, C., & Towers, G. J. (2021). SARS-CoV-2 sensing by RIG-I and MDA5 links epithelial infection to macrophage inflammation. The EMBO Journal, 40(15), e107826. https://doi.org/10.15252/embj.2021107826

Thurman, A. L., Ratcliff, J. A., Chimenti, M. S., & Pezzulo, A. A. (2021). Differential gene expression analysis for multi-subject single cell RNA sequencing studies with aggregateBioVar. Bioinformatics, 37(19), 3243–3251. https://doi.org/10.1093/bioinformatics/btab337

Triana, S., Metz-Zumaran, C., Ramirez, C., Kee, C., Doldan, P., Shahraz, M., Schraivogel, D., Gschwind, A. R., Sharma, A. K., Steinmetz, L. M., Herrmann, C., Alexandrov, T., Boulant, S., & Stanifer, M. L. (2021). Single-cell analyses reveal SARS-CoV-2 interference with intrinsic immune response in the human gut. Molecular Systems Biology, 17(4), e10232. https://doi.org/10.15252/msb.202110232

UniProt Consortium (2015). UniProt: a hub for protein information. Nucleic Acids Research, 43(Database issue), D204–D212. https://doi.org/10.1093/nar/gku989

Vallejos, C. A., Richardson, S., & Marioni, J. C. (2016). Beyond comparisons of means: understanding changes in gene expression at the single-cell level. Genome Biology, 17, 70. https://doi.org/10.1186/s13059-016-0930-3

Wilk, A. J., Rustagi, A., Zhao, N. Q., Roque, J., Martínez-Colón, G. J., McKechnie, J. L., Ivison, G. T., Ranganath, T., Vergara, R., Hollis, T., Simpson, L. J., Grant, P., Subramanian, A., Rogers, A. J., & Blish, C. A. (2020). A single-cell atlas of the peripheral immune response in patients with severe COVID-19. Nature Medicine, 26(7), 1070–1076. https://doi.org/10.1038/s41591-020-0944-y

Xia, H., Cao, Z., Xie, X., Zhang, X., Chen, J. Y., Wang, H., Menachery, V. D., Rajsbaum, R., & Shi, P. Y. (2020). Evasion of Type I Interferon by SARS-CoV-2. Cell Reports, 33(1), 108234. https://doi.org/10.1016/j.celrep.2020.108234

Xie, Y., & Ren, Y. (2019). Mechanisms of nuclear mRNA export: A structural perspective. Traffic, 20(11), 829–840. https://doi.org/10.1111/tra.12691

Yang, S. L., DeFalco, L., Anderson, D. E., Zhang, Y., Aw, J., Lim, S. Y., Lim, X. N., Tan, K. Y., Zhang, T., Chawla, T., Su, Y., Lezhava, A., Merits, A., Wang, L. F., Huber, R. G., & Wan, Y. (2021). Comprehensive mapping of SARS-CoV-2 interactions in vivo reveals functional virus-host interactions. Nature Communications, 12(1), 5113. https://doi.org/10.1038/s41467-021-25357-1

Yuan, S., Peng, L., Park, J. J., Hu, Y., Devarkar, S. C., Dong, M. B., Shen, Q., Wu, S., Chen, S., Lomakin, I. B., & Xiong, Y. (2020). Nonstructural protein 1 of SARS-CoV-2 is a potent pathogenicity factor redirecting host protein synthesis machinery toward viral RNA. Molecular Cell, 80(6), 1055–1066.e6. https://doi.org/10.1016/j.molcel.2020.10.034

Zanini, F., Robinson, M. L., Croote, D., Sahoo, M. K., Sanz, A. M., Ortiz-Lasso, E., Albornoz, L. L., Rosso, F., Montoya, J. G., Goo, L., Pinsky, B. A., Quake, S. R., & Einav, S. (2018). Virus-inclusive single-cell RNA sequencing reveals the molecular signature of progression to severe dengue. Proceedings of the National Academy of Sciences of the United States of America, 115(52), E12363–E12369. https://doi.org/10.1073/pnas.1813819115

Zhang, K., Miorin, L., Makio, T., Dehghan, I., Gao, S., Xie, Y., Zhong, H., Esparza, M., Kehrer, T., Kumar, A., Hobman, T. C., Ptak, C., Gao, B., Minna, J. D., Chen, Z., García-Sastre, A., Ren, Y., Wozniak, R. W., & Fontoura, B. (2021). Nsp1 protein of SARS-CoV-2 disrupts the mRNA export machinery to inhibit host gene expression. Science Advances, 7(6),eabe7386. https://doi.org/10.1126/sciadv.abe7386

Zheng, G. X., Terry, J. M., Belgrader, P., Ryvkin, P., Bent, Z. W., Wilson, R., Ziraldo, S. B., Wheeler, T. D., McDermott, G. P., Zhu, J., Gregory, M. T., Shuga, J., Montesclaros, L., Underwood, J. G., Masquelier, D. A., Nishimura, S. Y., Schnall-Levin, M., Wyatt, P. W., Hindson, C. M., Bharadwaj, R., … Bielas, J. H. (2017). Massively parallel digital transcriptional profiling of single cells. Nature Communications, 8, 14049. https://doi.org/10.1038/ncomms14049

Zhou, P., Yang, X. L., Wang, X. G., Hu, B., Zhang, L., Zhang, W., Si, H. R., Zhu, Y., Li, B., Huang, C. L., Chen, H. D., Chen, J., Luo, Y., Guo, H., Jiang, R. D., Liu, M. Q., Chen, Y., Shen, X. R., Wang, X., Zheng, X. S., … Shi, Z. L. (2020). A pneumonia outbreak associated with a new coronavirus of probable bat origin. Nature, 579(7798), 270–273. https://doi.org/10.1038/s41586-020-2012-7

Zhu, P., Lv, C., Fang, C., Peng, X., Sheng, H., Xiao, P., Kumar Ojha, N., Yan, Y., Liao, M., & Zhou, J. (2020). Heat shock protein member 8 is an attachment factor for infectious bronchitis virus. Frontiers in Microbiology, 11, 1630. https://doi.org/10.3389/fmicb.2020.01630

Ziegenhain, C., Vieth, B., Parekh, S., Reinius, B., Guillaumet-Adkins, A., Smets, M., Leonhardt, H., Heyn, H., Hellmann, I., & Enard, W. (2017). Comparative analysis of single-cell RNA sequencing methods. Molecular Cell, 65(4), 631–643.e4. https://doi.org/10.1016/j.molcel.2017.01.023

Ziegler, C., Allon, S. J., Nyquist, S. K., Mbano, I. M., Miao, V. N., Tzouanas, C. N., Cao, Y., Yousif, A. S., Bals, J., Hauser, B. M., Feldman, J., Muus, C., Wadsworth, M. H., 2nd, Kazer, S. W., Hughes, T. K., Doran, B., Gatter, G. J., Vukovic, M., Taliaferro, F., Mead, B. E., … HCA Lung Biological Network (2020). SARS-CoV-2 receptor ACE2 is an interferon-stimulated gene in human airway epithelial cells and is detected in specific cell subsets across tissues. Cell, 181(5), 1016–1035.e19. https://doi.org/10.1016/j.cell.2020.04.035

Zimmerman, K. D., Espeland, M. A., & Langefeld, C. D. (2021). A practical solution to pseudoreplication bias in single-cell studies. Nature Communications, 12(1), 738. https://doi.org/10.1038/s41467-021-21038-1

